# Heterogeneity in PHGDH protein expression potentiates cancer cell dissemination and metastasis

**DOI:** 10.1101/2021.01.24.427949

**Authors:** Matteo Rossi, Ginevra Doglioni, Laura Bornes, Dorien Broekaert, Mélanie Planque, Juan Fernández-García, Gianmarco Rinaldi, Joke Van Elsen, David Nittner, Cristina Jauset, Francesca Rizzollo, Carla Riera Domingo, Martin F Orth, Lacey E Dobrolecki, Thomas Van Brussel, Shao Thing Teoh, Arin B Aurora, Guy Eelen, Panagiotis Karras, Karl Sotlar, Harald Bartsch, Jean-Christophe Marine, Peter Carmeliet, Sean J Morrison, Michael T Lewis, Gregory J Hannon, Massimiliano Mazzone, Diether Lambrechts, Jacco van Rheenen, Thomas G P Grünewald, Sophia Y Lunt, Sarah-Maria Fendt

**Affiliations:** Laboratory of Cellular Metabolism and Metabolic Regulation, VIB-KU Leuven Center for Cancer Biology, VIB, Herestraat 49, 3000 Leuven, Belgium; Laboratory of Cellular Metabolism and Metabolic Regulation, Department of Oncology, KU Leuven and Leuven Cancer Institute (LKI), Herestraat 49, 3000 Leuven, Belgium; Division of Molecular Pathology, Oncode Institute, The Netherlands Cancer Institute, Amsterdam, The Netherlands; Histopathology Expertise Center, VIB-KU Leuven Center for Cancer Biology, 3000 Leuven, Belgium; Department of Oncology, KU Leuven, 3000 Leuven, Belgium; Cancer Research UK Cambridge Institute, Li Ka Shing Centre, University of Cambridge, Cambridge CB2 0RE, United Kingdom; Laboratory of Tumor Inflammation and Angiogenesis, Center for Cancer Biology (CCB), VIB, Leuven, Belgium; Laboratory of Tumor Inflammation and Angiogenesis, Department of Oncology, KU Leuven, Leuven, Belgium; Max-Eder Research Group for Pediatric Sarcoma Biology, Institute of Pathology, Faculty of Medicine, LMU Munich, Thalkirchner Strasse 36, 80337 Munich, Germany; StemMed, Ltd, Houston, TX, USA; Laboratory for Translational Genetics, VIB-KU Leuven Center for Cancer Biology, VIB, Leuven, Belgium; Laboratory for Translational Genetics, Department of Human Genetics, KU Leuven, Leuven, Belgium; Department of Biochemistry and Molecular Biology, Michigan State University, East Lansing, MI, USA; Children’s Research Institute and Department of Pediatrics, University of Texas Southwestern Medical Center, Dallas, TX 75390, USA; Laboratory of Angiogenesis and Vascular Metabolism, Department of Oncology, KU Leuven, Leuven, Belgium; Laboratory of Angiogenesis and Vascular Metabolism, Center of Cancer Biology, VIB, Leuven, Belgium; Laboratory of Molecular Cancer Biology, VIB Center for Cancer Biology, Leuven, Belgium; Department of Oncology, KULeuven, Leuven, Belgium; Department of Pathology, Paracelsus Medical University, SALK, Müllner Hauptstrasse 48, 5020 Salzburg, Austria; Institute of Pathology, Ludwig-Maximilians-University, 80337 Munich, Germany; Howard Hughes Medical Institute, University of Texas Southwestern Medical Center, Dallas, TX 75390, USA; Department of Molecular Biotechnology and Health Science, Molecular Biotechnology Centre, University of Torino, Torino, Italy; Hopp Children’s Cancer Center (KiTZ), Im Neuenheimer Feld 280, 69120 Heidelberg, Germany; Division of Translational Pediatric Sarcoma Research, German Cancer Research Center (DKFZ), German Cancer Consortium (DKTK), Im Neuenheimer Feld 280, 69120 Heidelberg, Germany; Institute of Pathology, Heidelberg University Hospital, Im Neuenheimer Feld 224, 69120 Heidelberg, Germany; Department of Chemical Engineering and Materials Science, Michigan State University, East Lansing, MI, USA

## Abstract

Cancer metastasis requires the transient activation of cellular programs enabling dissemination and seeding in distant organs. Genetic, transcriptional and translational intra-tumor heterogeneity contributes to this dynamic process. Beyond this, metabolic intra-tumor heterogeneity has also been observed, yet its role for cancer progression remains largely elusive. Here, we discovered that intra-tumor heterogeneity in phosphoglycerate dehydrogenase (PHGDH) protein expression drives breast cancer cell dissemination and metastasis formation. Specifically, we observed intra-tumor heterogeneous PHGDH expression in primary breast tumors, with low PHGDH expression being indicative of metastasis in patients. In mice, Phgdh protein, but not mRNA, expression is low in circulating tumor cells and early metastatic lesions, leading to increased dissemination and metastasis formation. Mechanistically, low PHGDH protein expression induces an imbalance in glycolysis that can activate sialic acid synthesis. Consequently, cancer cells undergo a partial EMT and show increased p38 as well as SRC phosphorylation, which activate cellular programs of dissemination. In turn, inhibition of sialic acid synthesis through knock-out of cytidine monophosphate N-acetylneuraminic acid synthetase (CMAS) counteracts the increased cancer cell dissemination and metastasis induced by low PHGDH expression. In conclusion, we find that heterogeneity in PHGDH protein expression promotes cancer cell dissemination and metastasis formation.

## INTRODUCTION

Inter- and intra-tumor heterogeneity are important determinants of metastatic potential and cancer progression^1–3^. Inter-tumor heterogeneity is the basis of classifying cancer subtypes, tumor grading, and selecting specific treatments^4^. Metabolically, inter-tumor heterogeneity has been widely studied, showing, for example, that *in vivo* clear cell renal carcinomas exhibit a glycolytic metabolism^5^, whereas human non-small-cell lung cancers subdivide into tumors that do or do not use lactate as a carbon source^6^. Mechanistically, inter-tumor metabolic heterogeneity can result from (epi)genetic alterations, the origin of cancer cells as well as the microenvironment and leads to the differential activation and dependence on various metabolic pathways^7–10^. With respect to tumor progression, it has been shown that inter-tumor heterogeneity in lactate utilization through monocarboxylate transporter 1 (MCT1) defines the metastatic potential of primary melanoma^11^. Thus, inter-tumor heterogeneity at all levels, including metabolism, has been reported.

Intra-tumor heterogeneity is closely associated with cancer progression and minimal residual disease leading to tumor relapse^1, 12^. Clonal heterogeneity, based on genetic alterations in subpopulations of cancer cells within a primary tumor, can contribute, through fitness selection, to metastatic progression and therapy resistance in minimal residual disease^1, 12, 13^. Nonclonal heterogeneity at the transcriptional and translational level promotes the plasticity of the cancer cells by allowing the transient activation of cellular programs^14, 15^. These, in turn, enable the cancer cells to dynamically change their phenotypical state and transition through the different steps of the metastatic cascade^16^. Indeed, tumor subpopulations with different degrees of epithelial-to-mesenchymal transition (EMT) had similar tumor-propagating capacities, but greatly varied in their potential to invade and metastasize, due to their differential ability to adopt a plastic behavior^17^. In particular, cancer cells that undergo a partial EMT have been identified to be highly competent in initiating metastasis formation^18^. While intra-tumor heterogeneity can result from nutrient and oxygen availability^19–22^, little is known about how heterogeneity and plasticity in the expression of metabolic enzymes within primary tumors and throughout the metastatic process contribute to cancer progression.

Here, we investigate intra-tumor heterogeneity in phosphoglycerate dehydrogenase (PHGDH) expression in primary tumors and plasticity in its overall expression during the metastatic process. We find that heterogeneous PHGDH expression in primary tumors promotes metastatic dissemination, and that dynamic modulations of PHGDH protein, but not mRNA, expression contributes to the plastic behavior of metastasizing cancer cells.

## RESULTS

### Heterogeneous and low PHGDH protein expression in primary tumors is indicative of metastasis

PHGDH is the first enzyme of the serine biosynthesis pathway, in which it converts 3-phosphoglycerate to 3-phosphonooxypyruvate. PHGDH is overexpressed and/or amplified in 70% of triple-negative breast cancers (TNBC) and some other cancers such as melanoma^23, 24^ and its activity is important for cancer proliferation^25–28^. However, to which extent intra-tumor heterogeneity in PHGDH protein expression exists remains elusive. We selected 129 predominately grade 2 and 3 invasive ductal carcinomas of the breast that were treated by primary surgical resection and were previously collected (1988-2006)^29^. We analyzed PHGDH protein expression in these primary breast cancers using immunohistochemistry. Subsequently, the PHGDH staining was evaluated by a pathologist for heterogeneity and intensity (**Extended Data Table 1**). We observed that 67% (86 out of 129) of the analyzed tumors from human breast cancer patients showed a homogeneous high PHGDH expression, while 33% showed heterogeneous (30 out of 129) or low (13 out of 129) PHGDH expression (**Figure 1a**). Next, we asked whether homogeneous versus heterogeneous/low PHGDH expression correlated with clinical parameters of cancer progression. Homogeneous versus heterogeneous/low PHGDH expressing tumors were similarly distributed across the different tumor grades (**Figure 1b**). In line with the known effect of PHGDH on cancer proliferation, tumor stage (pT) was higher in homogeneous versus heterogeneous/low PHGDH-expressing tumors (**Figure 1c**). Unexpectedly however, lymph node stage (pN) was significantly more advanced in heterogeneous/low PHGDH-expressing tumors compared to homogeneous PHGDH-expressing tumors (**Figure 1d**). Accordingly, approximately 60% more patients with heterogeneous/low PHGDH-expressing tumors showed distant metastases (14 out of 43; **Figure 1e**) compared to patients with homogeneous PHGDH-expressing tumors (17 out of 86; **Figure 1e**). Thus, we concluded that heterogeneous and low PHGDH protein expression is indicative of metastasis in triple negative breast cancer patients.

**Figure 1:**
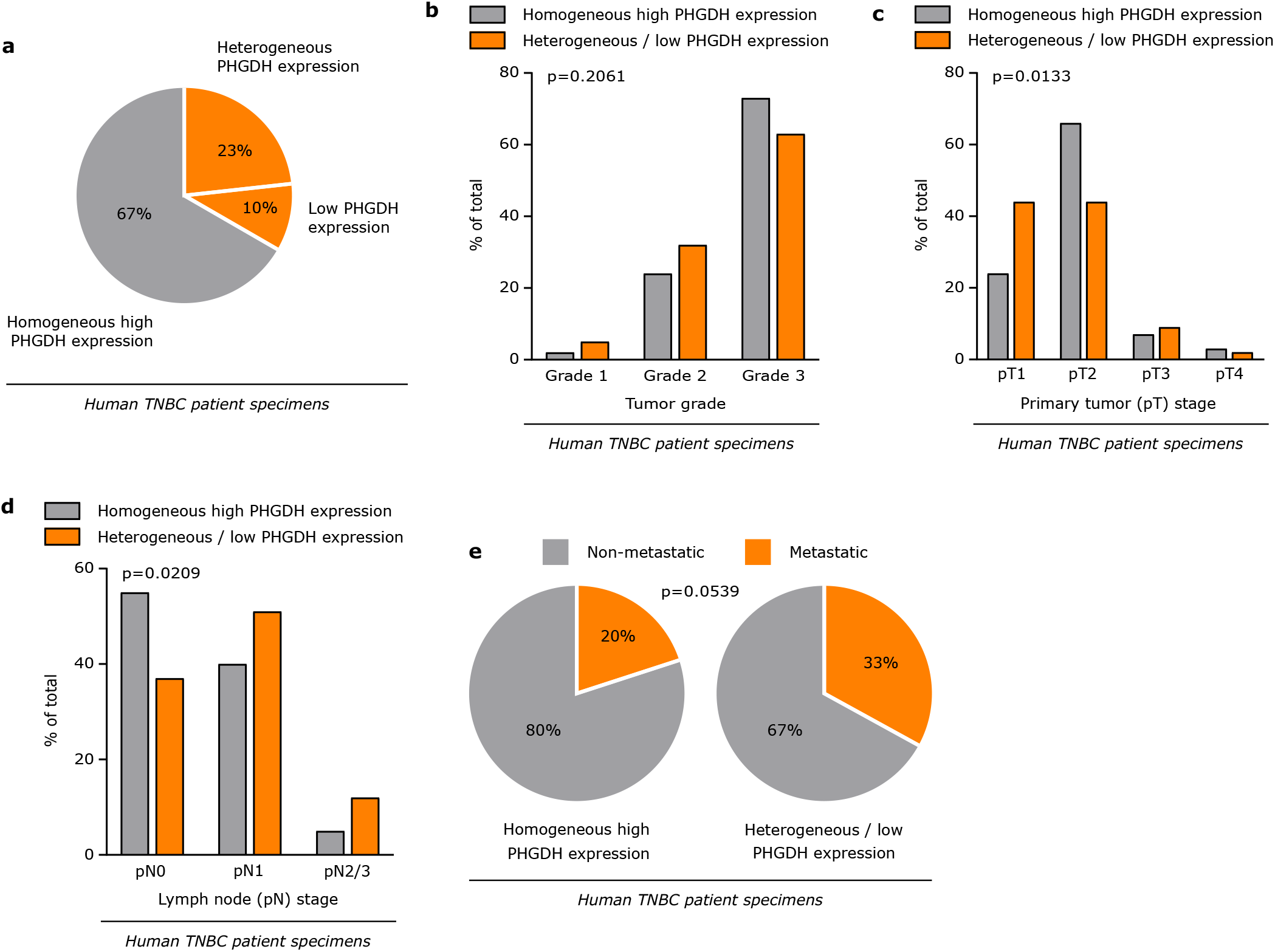
Heterogeneous and low PHGDH protein expression in primary tumors is indicative of metastasis. **a.** Distribution of PHGDH expression in human TNBC primary tumor specimens. PHGDH expression was assessed by immunohistochemistry. **b.** Comparison of the tumor grade in human TNBC primary tumors with homogeneous high and heterogeneous/low PHGDH expression. Chi-squared test. **c.** Comparison of the tumor stage (pT) in human TNBC primary tumors with homogeneous high and heterogeneous/low PHGDH expression. Chi-squared test. **d.** Comparison of the lymph node stage (pN) in human TNBC primary tumors with homogeneous high and heterogeneous/low PHGDH expression. Chi-squared test. **e.** Metastasis occurrence in TNBC patients bearing primary tumors with homogeneous high (17 out of 86) or heterogeneous/low (14 out of 43) PHGDH expression. Fisher’s exact test, two-sided. Total number of patients in the cohort, n=129; with homogeneous high expression, n=86; with heterogeneous expression, n=30; with low expression, n=13.

### Circulating tumor cells and early metastatic lesions exhibit low PHGDH expression

To further investigate the identified PHGDH heterogeneity in human breast tumors, we relied on mouse models. Similar to human invasive ductal carcinomas, mouse primary 4T1- and PDX-derived (BCM-5471) breast tumors displayed intra-tumor heterogeneity in PHGDH protein expression (**Figure 2a, 2b**). PHGDH-low 4T1 and PDX tumor cells within the primary breast tumor were in a slow cycling state, as evident from the positive correlation between the expression of PHGDH and the proliferation markers Ki67 and phospho-histone H3 (PHH3), respectively, based on imaging mass cytometry and immunohistochemistry (**Figure 2c, 2d**). Interestingly, slow cycling tumor cells have previously been associated with metastasis initiation capacity^22^. Therefore, we asked whether average PHGDH expression changes during metastatic progression. To address this question, we analyzed PHGDH protein expression in circulating tumor cells of three triple-negative breast cancer PDX mouse models (BCM-3107-R2TG18, BCM-3611-R3TG4 and BCM-4272-R3TG6) and compared it to the expression in the corresponding primary tumors. Strikingly, circulating tumor cells from all three PDX-models showed a 2.5- to 10.7-fold decrease in median PHGDH protein expression compared to the corresponding orthotopic primary tumors (**Figure 2e, Extended Data Figure 1a**). This result demonstrates that during metastatic progression, i.e. in the circulation, disseminating breast cancer cells are characterized by a low level of PHGDH protein expression.

**Figure 2:**
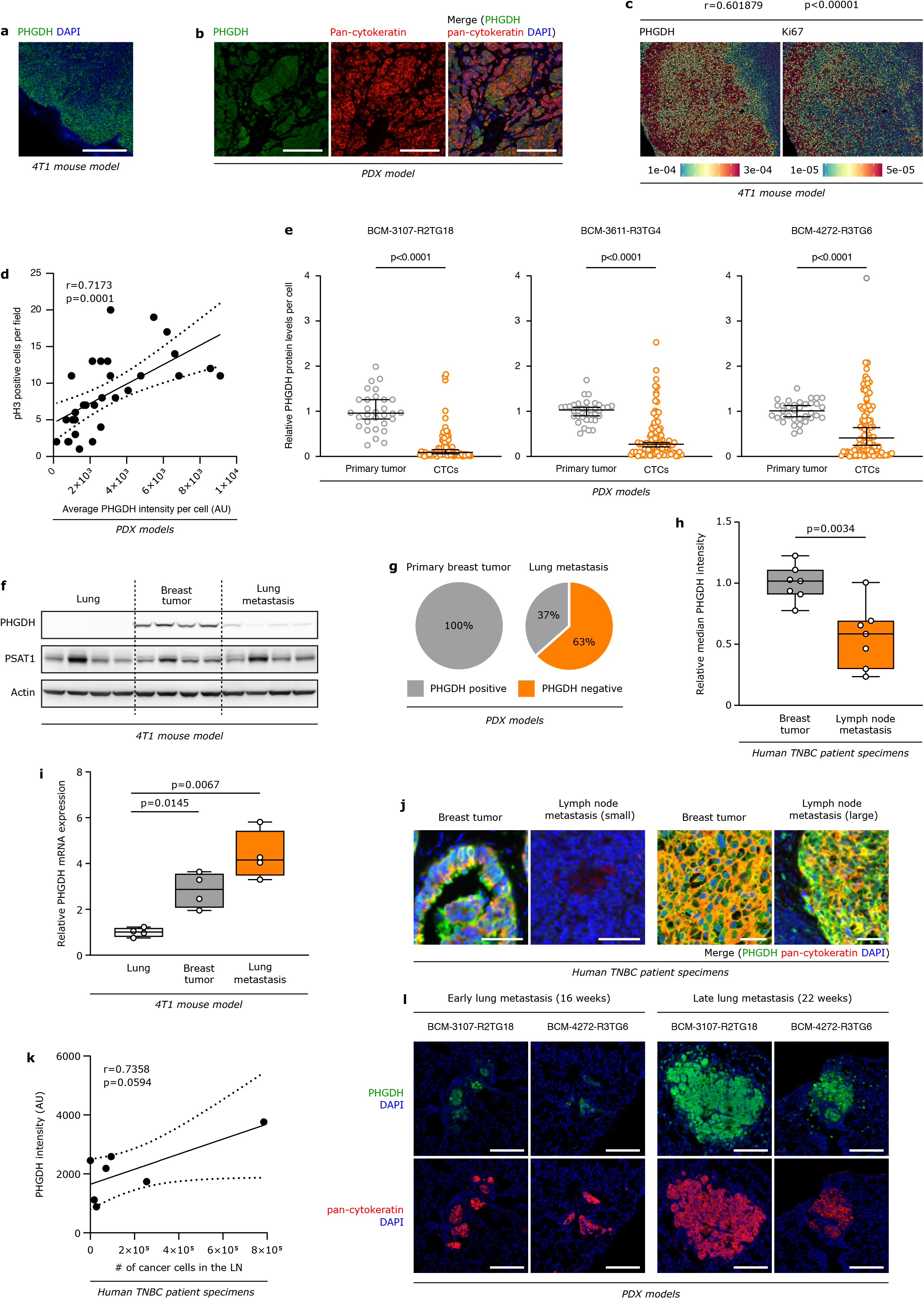
Circulating tumor cells and early metastatic lesions exhibit low PHGDH expression. **a.** Representative picture of Phgdh protein heterogeneity in the primary tumor from orthotopic (mammary fat pad, m.f.p.) 4T1 mouse model, assessed by immunohistochemistry. Green, Phgdh; blue, DAPI nuclear staining. Scale bar 1 mm. **b.** Representative picture of Phgdh protein heterogeneity in the primary tumor from orthotopic (m.f.p.) TNBC PDX model, assessed by immunohistochemistry. Green, Phgdh; red, pan-cytokeratin tumor marker; blue, DAPI nuclear staining. Scale bar 200 μm. **c.** Distribution and correlation of Phgdh and Ki67 protein expression in the primary tumor from orthotopic (m.f.p.) 4T1 mouse model, assessed by imaging mass cytometry. Nonparametric Spearman rank correlation. **d.** Correlation of PHGDH and phospho-histone H3 (PHH3) protein expression in the primary tumor from orthotopic (m.f.p.) TNBC PDX model, assessed by immunohistochemistry. The pooled analysis of 3 different PDX models, on 9 randomly chosen microscopy fields for each model, is shown in the graph. Nonparametric Spearman rank correlation. **e.** Expression levels of PHGDH protein in circulating tumor cells (CTCs) compared to the respective primary tumors from orthotopic (m.f.p.) TNBC PDX models, assessed by immunohistochemistry. Analysis performed on PDX models BCM-3107-R2TG18 (6 mice, 30 randomly chosen microscopy fields for the primary tumors, 5 per mouse, 87 single CTCs), BCM-3611-R3TG4 (7 mice, 35 randomly chosen microscopy fields for the primary tumors, 5 per mouse, 104 single CTCs) and BCM-4272-R3TG6 (7 mice, 35 randomly chosen microscopy fields for the primary tumors, 5 per mouse, 101 single CTCs). The solid lines indicate the median, the whiskers indicate the 95% confidence interval. Unpaired t test with Welch’s correction, two-tailed. **f.** Western blot analysis of Phgdh and Psat1 in lungs, primary breast tumors and lung metastases from orthotopic (m.f.p.) 4T1 mouse model (n=4). **g.** Positivity to PHGDH in primary breast tumors (n=9) and early lung metastases (~16 weeks after primary tumor initiation; n=52) from orthotopic (m.f.p.) TNBC PDX models, assessed by immunohistochemistry. **h.** Expression levels of PHGDH protein in lymph node metastases and matching primary breast tumors from TNBC patients (n=7), assessed by immunohistochemistry. The solid lines indicate the median, the boxes extend to the 25th and 75th percentiles, the whiskers span the smallest and the largest values. Unpaired t test with Welch’s correction, two-tailed. **i.** Relative change in *Phgdh* gene expression in lungs, primary breast tumors and lung metastases from orthotopic (m.f.p.) 4T1 mouse model (n=4). **j.** Representative pictures of PHGDH protein expression in small and large lymph node metastases and matching primary breast tumors from TNBC patients, assessed by immunohistochemistry. Green, PHGDH; red, pan-cytokeratin tumor marker; blue, DAPI nuclear staining. Scale bar 50 μm. **k.** Correlation of PHGDH protein expression with the lymph node metastasis size, determined by the number of cancer cells present in the lesion, in lymph node from TNBC patients (n=7), assessed by immunohistochemistry. Pearson correlation. **l.** Representative pictures of PHGDH protein expression in early (16 weeks) and late (22 weeks) lung metastases from orthotopic (m.f.p.) TNBC PDX models, assessed by immunohistochemistry. Green, PHGDH; red, pan-cytokeratin tumor marker; blue, DAPI nuclear staining. Scale bar 200 μm.

Next, we investigated average PHGDH protein expression in early (small) metastatic lesions and compared them to the corresponding primary breast tumors in mouse models and human samples. While primary tumors of 4T1 breast cancer cells showed heterogeneous but on average high Phgdh protein expression, early lung metastases were defined by low average Phgdh protein expression (**Figure 2f**). Comparable results were obtained with a second syngeneic and orthotopic breast cancer mouse model (EMT6.5; **Extended Data Figure 1b**) and with two different PDX-melanoma mouse models (UM12, M481; **Extended Data Figure 1c**). Similarly, 63% (33 out of 52) of early lung metastases from the three different PDX-mammary carcinoma mouse models exhibited average low or no PHGDH protein expression compared to the corresponding primary tumors (**Figure 2g**). Consistently, lymph node metastases from patients with breast cancer **(Extended Data Table 2)** showed a lower average PHGDH protein expression compared to the matching primary breast tumors (**Figure 2h, Extended Data Figure 1d**). Subsequently, we sought to relate this finding to altered *Phgdh* mRNA expression. Unexpectedly, however, we discovered that *Phgdh* mRNA expression did not decrease in early 4T1 and EMT6.5 metastases compared to the corresponding primary tumors (**Figure 2i, Extended Data Figure 1e**). Thus, we concluded that breast cancer cells express on average low levels of PHGDH protein, despite maintaining mRNA expression, during dissemination and metastasis formation.

Subsequently, we asked whether PHGDH protein expression was related to metastasis size. Indeed, we observed a correlation between PHGDH protein expression and lung metastasis size in the 4T1 breast cancer mouse model (**Extended Data Figure 1f**) and with lymph node metastases in human breast cancer patients (**Figure 2j, 2k**). These data suggest that PHGDH expression correlates with metastases size and, potentially, proliferation. To address this possibility, we analyzed PHGDH protein expression in lung metastases at two time points in two different PDX mouse models (BCM-3107-R2TG18 and BCM-4272-R3TG6). Remarkably, we discovered that early metastatic lesions (16 weeks after primary tumor initiation) had lower PHGDH protein expression compared to advanced (late; 22 weeks after primary tumor initiation) metastatic lesions (**Figure 2l, Extended Data Figure 1g**). These data are consistent with the notion that PHGDH protein expression is maintained at low levels during dissemination and re-acquired when metastases establish. This finding is in line with our recent study showing that PHGDH-mediated serine biosynthesis boosts mTORC1 signaling, promoting metastatic outgrowth^25^. Based on these data, we concluded that PHGDH protein expression is low in circulating tumor cells as well as early metastatic lesions and re-gained in late metastatic lesions of mouse models and human patients.

### Low PHGDH expression drives breast cancer cell dissemination and seeding in vivo

We then asked whether loss of PHGDH expression increases cancer cell dissemination and lung metastasis. To address this question, we silenced *Phgdh* in 4T1 breast cancer cells and performed time-lapse intravital imaging^30, 31^ (**Figure 3a**). Specifically, we expressed the fluorescent proteins mTurquoise in control cells and Dendra in *Phgdh*-silenced 4T1 breast cancer cells. Subsequently, a mix of control and *Phgdh*-silenced 4T1 breast cancer cells was injected into the mammary fat pad of mice. Upon reaching an approximate volume of 200 mm^3^, the tumors were surgically exposed and intravital images were acquired. Within each imaging field, the migratory behavior of the same number of randomly picked control and *Phgdh*-silenced 4T1 breast cancer cells was assessed. Using this *in vivo* analysis, we found a significantly higher number of migratory cells in the 4T1-silenced population than in the control population (**Figure 3b**). Furthermore, within the migratory population (>4 μm/h), *Phgdh*-silenced cells demonstrated higher displacement compared to control cells (**Figure 3c, Extended Data Video 1 and 2**). This finding demonstrates that loss of PHGDH expression drives *in vivo* cell migration, which is the first step of dissemination from primary tumors. Based on this finding, we expected that loss of Phgdh resulted in an increased number of cancer cells seeding in the lung. Thus, we analyzed the number of early metastatic lung lesions upon *Phgdh* silencing in 4T1 and EMT6.5 breast cancer mouse models. In line with the data from intravital imaging, we observed that *Phgdh* silencing in primary tumors increased the number of early metastatic lung lesions by 4.5-fold and 1.7-fold compared to control, respectively (**Figure 3d**). Based on these findings, we concluded that PHGDH loss in primary breast cancers potentiates *in vivo* dissemination and early metastatic seeding of cancer cells in the lungs of mice.

**Figure 3:**
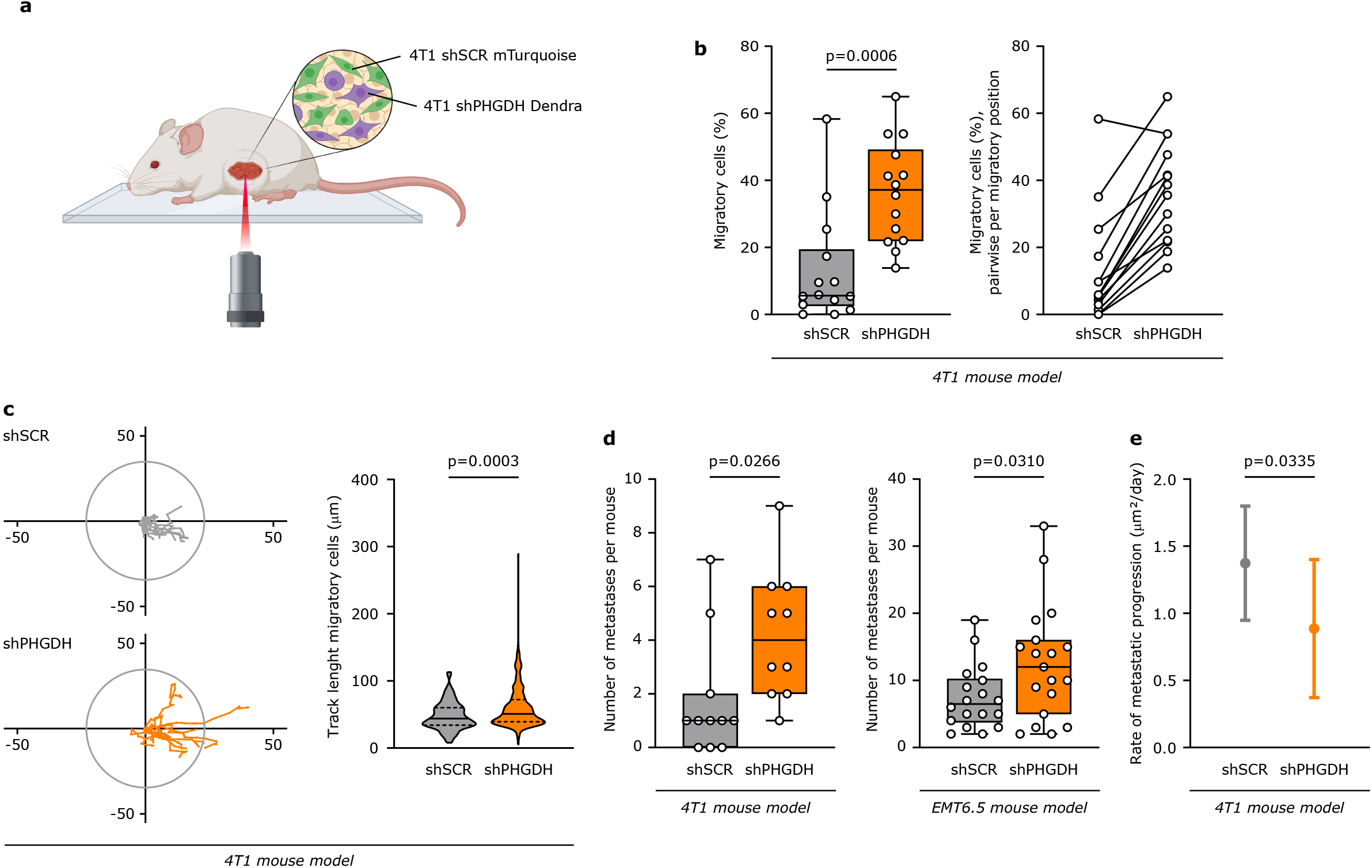
Low PHGDH expression drives breast cancer cell dissemination and seeding in vivo. **a.** Schematic representation of the time-lapse intravital imaging experiment setup. **b.** Percentage of migratory cells per migratory position (n=14) in the primary tumor from orthotopic (m.f.p.) 4T1 mouse models, assessed by time-lapse intravital imaging (n=8). Mice were injected with a mixture of 4T1 shSCR mTurquoise or 4T1 shPHGDH Dendra. The solid lines indicate the median, the boxes extend to the 25th and 75th percentiles, the whiskers span the smallest and the largest values. Unpaired t test with Welch’s correction, two-tailed. **c.** Track length of migratory cells in the primary tumor from orthotopic (m.f.p.) 4T1 mouse models, assessed by time-lapse intravital imaging. *Left panel*, representative tracks for 4T1 shSCR mTurquoise (n=9) and 4T1 shPHGDH Dendra (n=10). *Right panel*, track length of migratory 4T1 shSCR mTurquoise and 4T1 shPHGDH Dendra. The solid lines indicate the median, the dashed lines indicate the 25th and 75th percentiles. Unpaired t test with Welch’s correction, two-tailed. **d.** *Left panel*, number of lung metastases per mouse in orthotopic (m.f.p.) 4T1 mouse models, injected with either 4T1 shSCR (n=11) or 4T1 shPHGDH cells (n=10). *Right panel*, number of lung metastases per mouse in orthotopic (m.f.p.) EMT6.5 mouse models, injected with either EMT6.5 shSCR (n=18, two cohorts) or EMT6.5 shPHGDH cells (n=19, two cohorts). The solid lines indicate the median, the boxes extend to the 25th and 75th percentiles, the whiskers span the smallest and the largest values. Unpaired t test with Welch’s correction, two-tailed. **e.** Rate of metastatic progression of lung metastases from mice injected with either 4T1 shSCR (n=10 per time point) or 4T1 shPHGDH cells (n=10 per time point). Error bars represent standard deviation (s.d.) from mean. Unpaired t test with Welch’s correction, two-tailed.

Subsequently, we addressed the question whether low PHGDH expression affects metastatic outgrowth. The above-described data show that PHGDH expression is correlated with markers of proliferation (**Figure 2c, 2d**) and metastasis size (**Figure 2j, 2k, 2l, Extended Data Figure 1e, 1f**). Thus, we expect that forced low PHGDH expression may hinder metastatic outgrowth despite driving dissemination and early metastatic seeding. To address this possibility, we determined the rate of metastatic progression (area increase over time) of 4T1 lung metastases with and without *Phgdh* silencing. In line with our expectation, we observed that lung metastases with forced low Phgdh expression displayed a decreased rate of metastatic progression compared to control metastases that can regain Phgdh expression once successfully disseminated and seeded in the lung (**Figure 3e**). Taken together, we found that forced low Phgdh expression through *Phgdh* silencing increased *in vivo* dissemination and early metastatic seeding but hindered metastatic outgrowth. This finding is consistent with the notion that plasticity in PHGDH expression is functionally relevant for metastasis formation.

### Low PHGDH expression promotes sialic acid synthesis

To investigate the mechanism by which PHGDH loss drives dissemination and metastasis, we studied the metabolism of cultured 4T1 and EMT6.5 cancer cells upon *Phgdh* silencing, based on metabolomics analysis (**Extended Data Table 3**). We observed that the metabolite fructose bisphosphate was consistently depleted upon PHGDH silencing in both cancer cell lines (**Figure 4a**). The pentose phosphate pathway and sialic acid metabolism, which both have already been linked to metastasis^11, 32^, branch from glycolysis upstream of fructose bisphosphate (**Extended Data Figure 2a)**. Thus, we hypothesized that either of these pathways is induced by *Phgdh* silencing. To determine which pathway is of importance we performed RNA sequencing in cultured 4T1 cells upon *Phgdh* silencing compared to control. We observed no major changes in enzyme expression within the pentose phosphate pathway. However, the expression of several enzymes of the hexosamine and sialic acid metabolism was increased upon *Phgdh* silencing (**Figure 4b, Extended Data Table 4**), which included the key rate-limiting enzyme cytidine monophosphate N-acetylneuraminic acid synthetase (*Cmas*; **Figure 4c**). We confirmed the upregulation of *Cmas* in cultured 4T1 and EMT6.5 cancer cells upon PHGDH silencing and in metastases versus primary tumors in our 4T1 mouse model (**Figure 4d, 4e**). Based on these data we hypothesized that flux through the sialic acid pathway increased upon PHGDH loss. To test this hypothesis, we determined sialic acid metabolism activity in cultured 4T1 cancer cells upon *Phgdh* silencing based on dynamic ^13^C tracer analysis^33^. In line with the gene expression data, we observed an increased flux through the sialic acid pathway (**Figure 4f**) and increased levels of the main sialic acid pathway metabolites (**Extended Data Figure 2b**) upon *Phgdh* silencing in 4T1. Conversely, overexpression of PHGDH in MDA-MB-231 cells, which normally do not express PHGDH, led to decreased flux through the sialic acid pathway **(Extended Data Figure 2c**) and decreased levels of the sialic acid pathway metabolites (**Extended Data Figure 2d**). Based on these data, we concluded that low PHGDH expression activates sialic acid metabolism in breast cancer cells.

**Figure 4:**
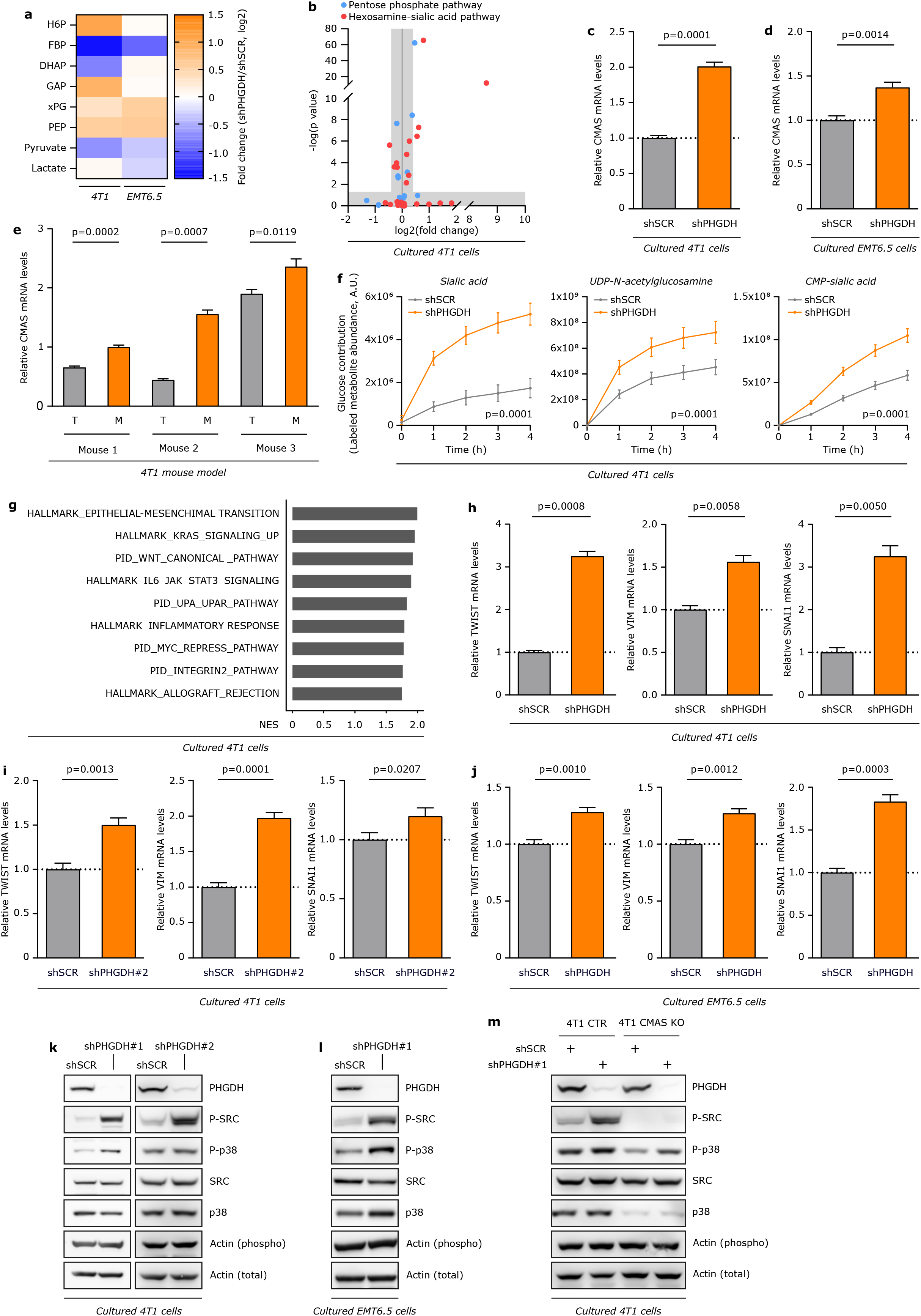
Low PHGDH expression promotes sialic acid synthesis and cellular programs downstream of sialic acid metabolism. **a.** Metabolite abundance changes upon *Phgdh* knock-down in 4T1 and EMT6.5 cells. Data represent fold changes compared to non-silenced cells (shSCR). H6P, hexose-6-phosphate; FBP, fructose bisphosphate; DHAP, dihydroxyacetone phosphate; GAP, glyceraldehyde 3-phosphate; xPG, 2/3-phosphoglycerate; PEP, phosphoenolpyruvate. **b.** Changes in the expression of genes in the pentose phosphate pathway (blue) and in the hexosamine and sialic acid pathway (red), extrapolated from bulk RNA sequencing analysis of 4T1 cells upon *Phgdh* knock-down (4T1 shPHGDH), compared to control cells (4T1 shSCR). The horizontal gray area defines p < 0.05; the vertical gray area defines fold changes ≤ 1.5. **c-d.** Relative change in *Cmas* gene expression upon *Phgdh* knock-down in 4T1 and EMT6.5 cells. **e.** Relative change in *Cmas* gene expression in primary tumors and matched lung metastases in orthotopic (m.f.p.) 4T1 mouse models. Error bars represent standard deviation (s.d.) from mean. **f.** Dynamic labeling of 4T1 cells, showing ^13^C glucose incorporation into sialic acid, UDP-N-acetylglucosamine and CMP-sialic acid upon *Phgdh* knock-down. Two-way ANOVA. Error bars represent standard deviation (s.d.) from mean. **g.** Gene Set Enrichment Analysis (GSEA) analysis of bulk RNA sequencing data from 4T1 cells upon *Phgdh* knock-down (4T1 shPHGDH), compared to control cells (4T1 shSCR). The highest-ranking Gene Ontology terms are depicted. **h-j.** Relative change in *Twist*, *vimentin* (*Vim*) and *Snai1* gene expression upon PHGDH knock-down in 4T1 and EMT6.5 cells. Error bars represent standard deviation (s.d.) from mean. **k-l.** Phosphorylation of proto-oncogene tyrosine-protein kinase Src (c-Src) and p38 mitogen-activated protein kinase upon *Phgdh* knock-down in 4T1 and EMT6.5 cells. **m.** Phosphorylation of proto-oncogene tyrosine-protein kinase Src (c-Src) and p38 mitogen-activated protein kinase upon *Phgdh* knock-down in 4T1 cells with functional (CTR) or inactive (*Cmas* KO) sialic acid pathway. Error bars represent s.d. from mean (n=3). For western blot and expression data all experiments were performed in triplicate, one representative experiment is shown. Unpaired t test with Welch’s correction, two-tailed, unless stated otherwise.

### Low PHGDH expression promotes a partial EMT, p38 and SRC phosphorylation

Next, we asked how low PHGDH expression promotes dissemination. To disseminate from the primary tumor cancer cells need to alter their cell-cell and cell-matrix interactions which can be achieved through alterations in glycosylation at the cell surface^34^. Moreover, at least in some tumors the induction of a partial EMT increases dissemination^17^. Hence, we assessed the expression of EMT markers and phosphorylation of p38 and of the proto-oncogene tyrosine-protein kinase Src (c-SRC), indicative of focal adhesion kinase (FAK) activation and thus of altered cell-cell and cell matrix interactions^35–38^, in 4T1 and EMT6.5 cancer cells. We identified EMT as the highest-ranking gene ontology term, along with other key pathways of migration, invasion and metastasis formation, based on RNA sequencing in 4T1 cells upon loss of Phgdh (**Figure 4g**). We validated these findings based on RT-PCR of molecular markers involved in these processes (**Extended Data Figure 2e**), among which the well-established EMT markers *vimentin*, *Twist* and *Snai1*, in *Phgdh*-silenced 4T1 and EMT6.5 cancer cells (**Figure 4h, 4i, 4j**). Moreover, p38 and c-Src phosphorylation were increased in *Phgdh*-silenced compared to control 4T1 and EMT6.5 cancer cells (**Figure 4k, 4l**). Subsequently, we asked whether overexpression of PHGDH in MDA-MB-231 breast cancer cells impaired the induction of these cellular programs. In accordance with our knockdown data, we found that PHGDH overexpression resulted in a decrease in p38 and c-SRC phosphorylation (**Extended Data Figure 2f**). We therefore concluded that PHGDH status modulates EMT marker expression and phosphorylation of p38 and c-SRC.

Next, we investigated whether the increase in sialic acid metabolism upon *Phgdh* silencing is responsible for the elevation of the identified cellular programs. We combined *Cmas* knockout, which will block the sialic acid pathway, with *Phgdh* silencing in 4T1 breast cancer cells, and evaluated the phosphorylation of p38 and c-Src. Strikingly, we observed that the combined Cmas and Phgdh loss decreased the phosphorylation of p38 and c-Src induced by *Phgdh* silencing (**Figure 4m**). Therefore, we concluded that loss of PHGDH results in a partial EMT and in an increase of p38 and c-SRC activation, mediated through sialic acid synthesis.

### Low PHGDH expression promotes sialic acid metabolism-mediated invasion and migration

We then examined whether PHGDH status and consequently sialic acid metabolism activity also modulated dissemination-associated cellular behavior. We measured spheroid and cell suspension invasion into a Matrigel-collagen I matrix, as well as transwell migration with and without coating of (lymphatic) endothelial cells. We found that 4T1 and EMT6.5 cancer cells with *Phgdh* silencing showed an increased ability to migrate through transwells pores as such or covered by a layer of (blood or lymphatic) endothelial cells (**Figure 5a, 5b**). Moreover, the invasive area and distance in Matrigel-collagen I were substantially elevated in *Phgdh*-silenced compared to control 4T1 and EMT6.5 cancer cells (**Figure 5c, 5d, 5e, 5f**). Subsequently, we asked whether PHGDH overexpression in MDA-MB-231 breast cancer cells impaired invasion. Consistent with our knockdown data, we observed that PHGDH overexpression resulted in a decreased invasive area in Matrigel-collagen I compared to control MDA-MB-231 breast cancer cells (**Figure 5g**). This demonstrates that PHGDH status modulates *in vitro* migration and invasion.

**Figure 5:**
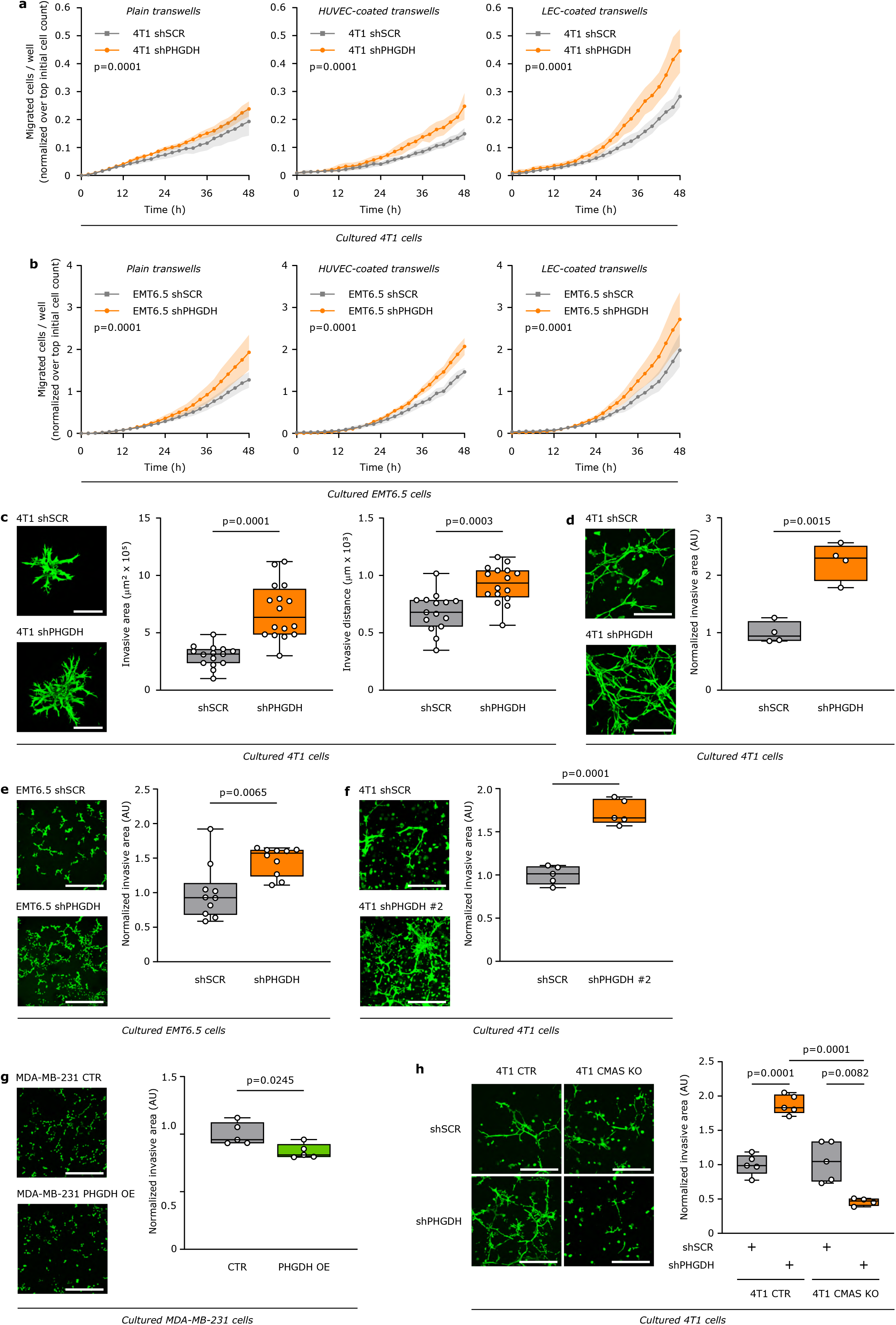
Low PHGDH expression promotes sialic acid metabolism-mediated invasion and migration. **a-b.** Migratory ability of 4T1 and EMT6.5 cells upon *Phgdh* knock-down (shPHGDH) compared to control (shSCR) cells. Migration was assessed in plain transwells and in transwells coated with either vascular endothelial cells (HUVECs) or lymphatic endothelial cells (LECs). Two-way ANOVA. **c-f.** Invasive ability in a 3D matrix of 4T1 and EMT6.5 cells upon *Phgdh* knock-down (shPHGDH) compared to control (shSCR) cells. Invasion was assessed by measuring the invasive area of cancer cells stained with calcein green. Representative images are depicted in the left panel (scale bar 500 μm), quantification in the right panel. Each dot represents a different, randomly selected microscopy field (n=15 for Figure 5c, n=5 for Figure 5d and Figure 5f, n=10 for Figure 5e). **g.** Invasive ability in a 3D matrix of MDA-MB-231 cells upon PHGDH overexpression (PHGDH OE) compared to control (CTR) cells. Invasion was assessed by measuring the invasive area of cancer cells stained with calcein green. Representative images are depicted in the left panel (scale bar 500 μm), quantification in the right panel. Each dot represents a different microscopy field (n=5). **h.** Invasive ability in a 3D matrix of 4T1 cells with functional (CTR) or inactive (*Cmas* KO) sialic acid pathway upon *Phgdh* knock-down (shPHGDH) compared to control (shSCR) cells. Invasion was assessed by measuring the invasive area of cancer cells stained with calcein green. Each dot represents a different microscopy field (n=5). Unpaired t test with Welch’s correction, two-tailed, unless otherwise stated. The solid lines indicate the median, the boxes extend to the 25th and 75th percentiles, the whiskers span the smallest and the largest values.

Subsequently, we investigated whether inhibition of sialic acid metabolism counteracted the low PHGDH expression-induced invasive phenotype. We assessed the invasion of 4T1 cancer cells with combined Cmas and Phgdh loss and found that *Cmas* knockout decreased the *Phgdh* silencing-induced invasive capacities of 4T1 cells in Matrigel-Collagen I (**Figure 5h**). Hence, we concluded that sialic acid synthesis mediates the low PHGDH expression-induced *in vitro* invasiveness.

### Low PHGDH protein expression, but not low catalytic activity, creates an imbalance in glycolysis

PHGDH catalyzes the conversion of 3-phosphoglycerate to 3-phosphonooxypyruvate which, in conjunction with two downstream enzymes, leads to the production of serine and α-ketoglutarate (**Figure 6a**). Thus, we asked whether the loss of PHGDH catalytic function or protein is required to activate sialic acid synthesis and downstream cellular programs of dissemination. In particular, we used the catalytic PHGDH inhibitor PH-755 (1 μM)^26, 39^, which reduced serine biosynthesis by 85% in 4T1 cancer cells (**Figure 6b**) without affecting protein expression (**Figure 6c)**. Next, we measured sialic acid metabolism flux upon PH-755 treatment of 4T1 cells based on dynamic ^13^C tracer analysis^33^. Unexpectedly, PH-755 treatment did not increase, but instead decreased, sialic acid metabolism flux (**Figure 6d**) and metabolite abundance (**Figure 6e**) in 4T1 cells. Substrate metabolite concentrations are important mediators in diverting flux into glycolytic side-pathways^40^. We found that *Phgdh* silencing (**Figure 4a**), but not catalytic inhibition with PH-755 (**Figure 6f**), substantially increased hexose-6-phosphate abundance and decreased fructose bisphosphate abundance. This metabolic alteration in upper glycolysis can support flux diversion towards sialic acid synthesis. The enzyme phosphofructokinase (PFK) competes with the sialic acid synthesis enzyme l-glutamine d-fructose 6-phosphate amidotransferase (GFAT) for fructose-6-phosphate. The three PFK isoforms (PFKL, PFKM and PFKP) are tightly regulated by post-translational modifications, allosteric modulation and protein-protein interactions^41, 42^. PFKM, in particular, is autocatalytic activated by fructose bisphosphate^43^ and thus inhibited by low fructose bisphosphate concentrations, while the expression of the enzyme aldolase, fructose-bisphosphate A (ALDOA) controls fructose bisphosphate concentration^44^. Strikingly, Phgdh showed a strong protein interaction with PfkM and PfkP (**Figure 6g**). Moreover, *Phgdh* silencing, but not catalytic inhibition with PH-755, increased the expression of AldoA (and PfkM) compared to control in 4T1 and EMT6.5 breast cancer cells (**Figure 6h**). Accordingly, lower glycolytic flux was increased upon *Phgdh* silencing, based on the assessment of tritiated H2O production at the enolase reaction downstream of AldoA (**Figure 6i**), while glucose uptake was not significantly altered (**Figure 6j**). These metabolic alterations demonstrate that low PHGDH protein expression induced an imbalance in glycolysis, which in turn could divert flux towards sialic acid synthesis.

**Figure 6:**
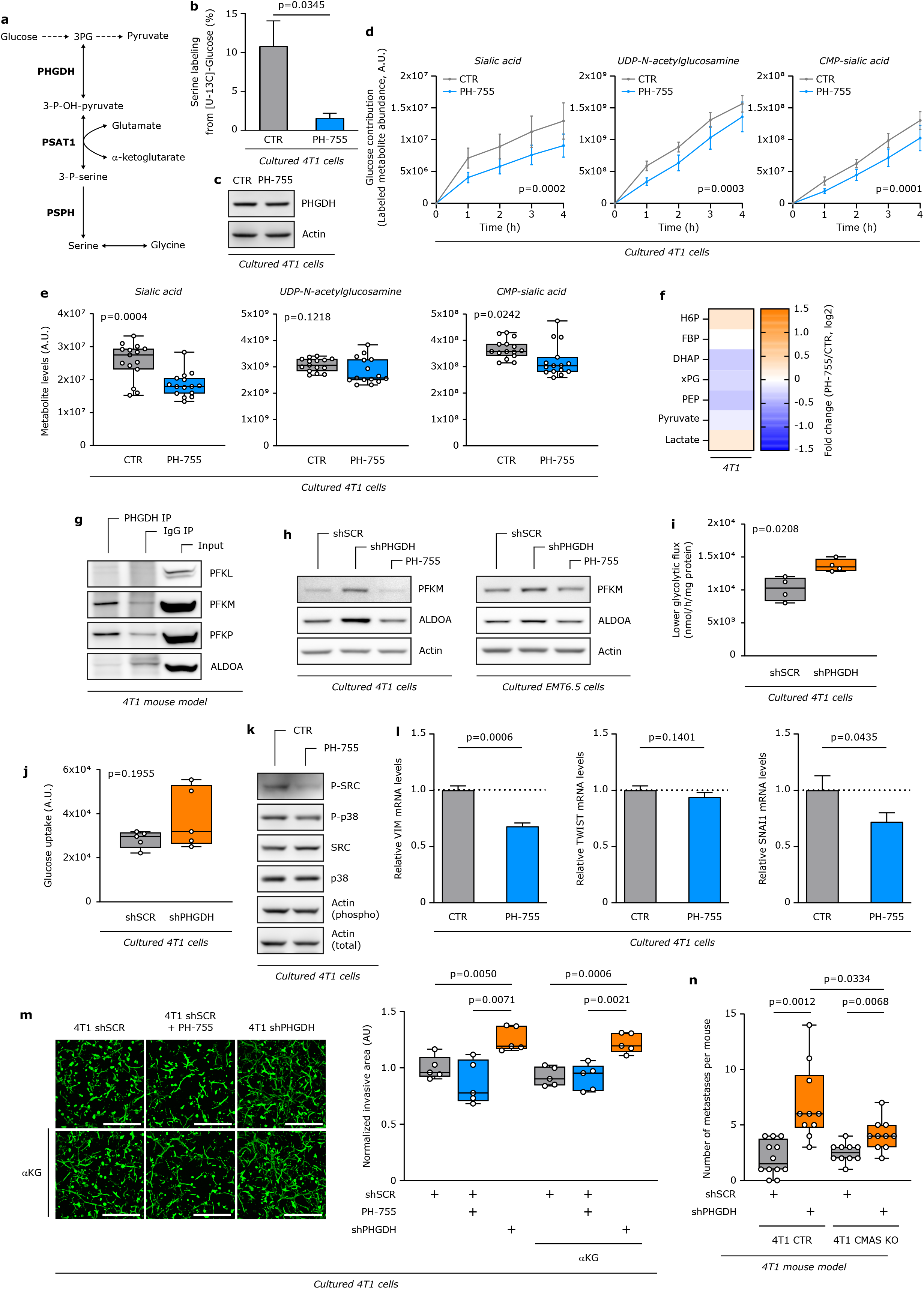
Low PHGDH protein expression creates an imbalance in glycolysis that promotes invasion and metastasis via the sialic acid pathway. **a.** Schematic representation of the *de novo* serine biosynthesis pathway branching from glycolysis at the level of 3PG. Enzymes are depicted in bold. Solid lines represent single reactions, dashed lines recapitulate multiple reactions. **b.** Inhibition of *de novo* serine biosynthesis assessed through measurement of serine m+3 labeling enrichment after incubation of 4T1 cells for 24h in culture medium containing ^13^C_6_ glucose, upon treatment with the PHGDH catalytic inhibitor PH-775 (1 μM) for 72h. Error bars represent s.d. from mean. **c.** Protein expression levels of PHGDH upon treatment with the PHGDH catalytic inhibitor PH-775 (1 μM) for 72h. **d.** Dynamic labeling of 4T1 cells, showing ^13^C glucose incorporation into sialic acid, UDP-N-acetylglucosamine and CMP-sialic acid upon treatment with the PHGDH catalytic inhibitor PH-775 (1 μM) for 72h. Two-way ANOVA. Error bars represent s.d. from mean. **e.** Metabolite abundances of sialic acid, UDP-N-acetylglucosamine and CMP-sialic acid upon treatment with the PHGDH catalytic inhibitor PH-775 (1 μM) for 72h in 4T1 cells. The solid lines indicate the median, the boxes extend to the 25th and 75th percentiles, the whiskers span the smallest and the largest values (n=15). Unpaired t test with Welch’s correction, two-tailed. **f.** Abundance changes in glycolytic intermediates upon PHGDH inhibition with PH-775 (1 μM) in 4T1 cells. Data represent fold changes compared to control cells. H6P, hexose-6-phosphate; FBP, fructose bisphosphate; DHAP, dihydroxyacetone phosphate; xPG, 2/3-phosphoglycerate; PEP, phosphoenolpyruvate. **g.** Interaction of Phgdh with glycolytic enzymes PfkL, PfkM, PfkP and Aldoa in the primary tumor of the orthotopic (m.f.p.) 4T1 mouse model. **h.** Protein expression levels of the glycolytic enzymes PfkM and Aldoa in 4T1 cells, upon *Phgdh* knock-down or *PHGDH* inhibition with PH-775 (1 μM). **i.** Lower glycolytic flux in 4T1 cells upon *Phgdh* knock-down (shPHGDH) compared to control (shSCR) cells based on radioactive H2O production by enolase. The solid lines indicate the median, the boxes extend to the 25th and 75th percentiles, the whiskers span the smallest and the largest values. Unpaired t test with Welch’s correction, two-tailed. **j.** Glucose uptake in 4T1 cells upon *Phgdh* knock-down (shPHGDH) compared to control (shSCR) cells. The solid lines indicate the median, the boxes extend to the 25th and 75th percentiles, the whiskers span the smallest and the largest values. Unpaired t test with Welch’s correction, two-tailed. **k.** Phosphorylation of proto-oncogene tyrosine-protein kinase Src (c-Src) and p38 mitogen-activated protein kinase in 4T1 cells upon treatment with the PHGDH catalytic inhibitor PH-775 (1 μM) for 72h. **l.** Relative change in *Twist*, *vimentin* (*Vim*) and *Snai1* gene expression in 4T1 cells upon treatment with the PHGDH catalytic inhibitor PH-775 (1 μM) for 72h. Error bars represent s.d. from mean. **m.** Invasive ability in a 3D matrix of 4T1 cells upon treatment with the PHGDH catalytic inhibitor PH-775 (1 μM) for 72h and rescue with cell-permeable α-ketoglutarate (αKG, 1 mM) where indicated. Invasion was assessed by measuring the invasive area of cancer cells stained with calcein green. Representative images are depicted in the left panel (scale bar 500 μm), quantification in the right panel. Each dot represents a different, randomly selected microscopy field. The solid lines indicate the median, the boxes extend to the 25th and 75th percentiles, the whiskers span the smallest and the largest values (n=5). **n.** Number of lung metastases per mouse in orthotopic (m.f.p.) 4T1 mouse models. Mice (n=10, except for 4T1 CTR shSCR, for which n=12) were injected with either 4T1 cells with functional (CTR) or inactive (*Cmas* KO) sialic acid pathway, upon *Phgdh* knock-down (shPHGDH) compared to control (shSCR) cells. The solid lines indicate the median, the boxes extend to the 25th and 75th percentiles, the whiskers span the smallest and the largest values. Unpaired t test with Welch’s correction, two-tailed. N=3 unless stated otherwise. For western blot and expression data all experiments were performed in triplicate, one representative experiment is shown. Unpaired t test with Welch’s correction, two-tailed, unless stated otherwise.

To provide functional relevance for the finding that only low PHGDH protein expression, but not loss catalytic activity, drives sialic acid synthesis, we analyzed cellular process of dissemination. Accordingly, PH-755 treatment did not increase p38 and c-Src phosphorylation (**Figure 6k**), nor did it elevate *vimentin*, *Twist* and *Snai1* expression (**Figure 6l**). Moreover, the invasion ability of 4T1 suspension cells in Matrigel was not altered by PH-755 treatment (**Figure 6m**). Similarly, α-ketoglutarate supplementation (1 mM) did not change the invasive area of 4T1 cells in Matrigel-Collagen I (**Figure 6n**). We therefore concluded that loss of PHGDH protein, but not catalytic function, created an imbalance in glycolysis that could activate sialic acid synthesis-mediated cellular programs of dissemination.

### Sialic acid metabolism mediates in vivo the metastatic seeding induced by low PHGDH protein levels

Finally, we hypothesized that inhibiting sialic acid metabolism through CMAS deletion counteracts the PHGDH low-induced *in vivo* metastatic seeding. To address this hypothesis, we injected 4T1 cells with combined Cmas and Phgdh loss and *Phgdh*-silenced 4T1 cells, as well as their respective controls, into the mammary fat pad of mice and we determined the number of early lung metastatic lesions. We observed that control 4T1 cells and sole *Cmas* knockout cells resulted in the same number of early lung metastatic lesions (**Figure 6l**), whereas, as expected, loss of Phgdh alone increased the number of early lung metastatic lesions (**Figure 6l**). Strikingly, and in line with our *in vitro* results, we further found that combined Cmas and Phgdh loss greatly reduced the number of early lung metastatic lesions almost to the same level as control 4T1 cells and sole *Cmas* knockout cells (**Figure 6l**). Based on these data, we concluded that sialic acid metabolism mechanistically links low PHGDH protein expression to *in vivo* metastatic dissemination.

## DISCUSSION

Here, we discovered that PHGDH protein expression is dynamically regulated during metastasis formation, with low PHGDH expression potentiating dissemination of cancer cells from the primary tumor via activation of sialic acid metabolism. Accordingly, heterogeneous and low PHGDH protein expression is associated with increased metastasis in breast cancer patients.

Metabolic rewiring has been linked to cancer metastasis, with increased activity of certain metabolic pathways often found in metastases compared to the primary tumor^45^. However, much less is understood about metabolic heterogeneity and the dynamic modulation of metabolic enzymes during the metastatic process^46^. Particularly, the importance of transiently downregulating the expression of certain enzymes to alter global cellular programs remains largely elusive.

There is an extensive body of literature showing that increased PHGDH activity is important for cancer cell proliferation^47^. In this respect it has been shown that PHGDH activity yielding serine is important for nucleotide synthesis, redox metabolism and one-carbon metabolism^45, 48–51^. Moreover, α-ketoglutarate, which is produced in equimolar amounts with serine, was shown to activate growth signaling through mTORC1 activation^25^. Accordingly, in low serine environments such as the brain and in environments that boost the activity of the serine biosynthesis pathway such as the lung, PHGDH activity is necessary for effective proliferation^26^ and growth signaling^25^. In line with this, average high PHGDH expression in multiple tumor types has been associated with poor prognosis^45, 47^.

Our research adds a new level of understanding to the biology of PHGDH by showing that heterogeneous rather than homogenous PHGDH expression is associated with increased metastasis in breast cancer patients. Indeed, PHGDH protein expression is dynamically regulated during breast and, potentially, melanoma metastasis, and its transient downregulation in the early steps of the metastatic process potentiates the dissemination of cancer cells from the primary tumor. Although the underlying mechanism is completely different, our finding is complementary to the observation that stem cells differentiate when *de novo* serine biosynthesis is activated^52^. Importantly, the mechanism identified here depends on a non-catalytic function of PHGDH. The moonlighting function of PHGDH is highly understudied, with only two additional reports suggesting that PHGDH expression stabilizes the pro-tumor transcription factor forkhead box M1 (FOXM1) in glioma cells^53^ and promotes proliferation through an interaction with translation initiation factors eIF4A1 and eIF4E in pancreatic cancer cells^54^. Strikingly we describe for the first time the importance of intra-tumor heterogeneous PHGDH protein expression in primary tumors and provide functional relevance for PHGDH downregulation for potentiating metastatic dissemination. Thus, our study may ignite further research in this new topic.

In conclusion, while PHGDH supports cancer cell proliferation through its metabolic function, low PHGDH protein expression can non-catalytically drive cancer dissemination and metastasis formation via sialic acid metabolism.

## AUTHOR CONTRIBUTIONS

Conception and design: MR and S-MF.

Development of methodology: MR, GD, DB, MP, DN, LB, CJ, and CRD.

Acquisition of data (including providing animals, acquiring and managing patient samples, providing facilities, etc.): MR, GD, DB, MP, GR, JVE, DN, LB, CJ, FR, MFO, LED, TVB, STT, ABA, GE, PK, KS, HB, J-CM, PC, SJM, MTL, GJH, MM, DL, JvR, TGPG and SYL.

Analysis and interpretation of data (e.g., statistical analysis, biostatistics, computational analysis): MR, GD, JF-G, LB, CJ, MFO, STT, JvR, TGPG and S-MF.

Writing, review, and/or revision of the manuscript: MR and S-MF.

Administrative, technical, or material support (i.e., reporting or organizing data, constructing databases): MR, GD, DB, MP and JVE.

Study supervision: S-MF.

## ACKNOWLEDGEMENTS

We would like to thank Matt Vander Heiden (MIT) for providing the PHGDH overexpression plasmid. We would like to thank Pawel Bieniasz-Krzywiec (VIB-KU Leuven) for his help with the transwell migration assay and Vincent van Hoef (VIB Bioinformatics Core Facility) for his help with the RNAseq analysis. We would like to thank Raze Therapeutics for providing us with the PHGDH inhibitor PH-755.

---

**Extended Data Figure 1:**
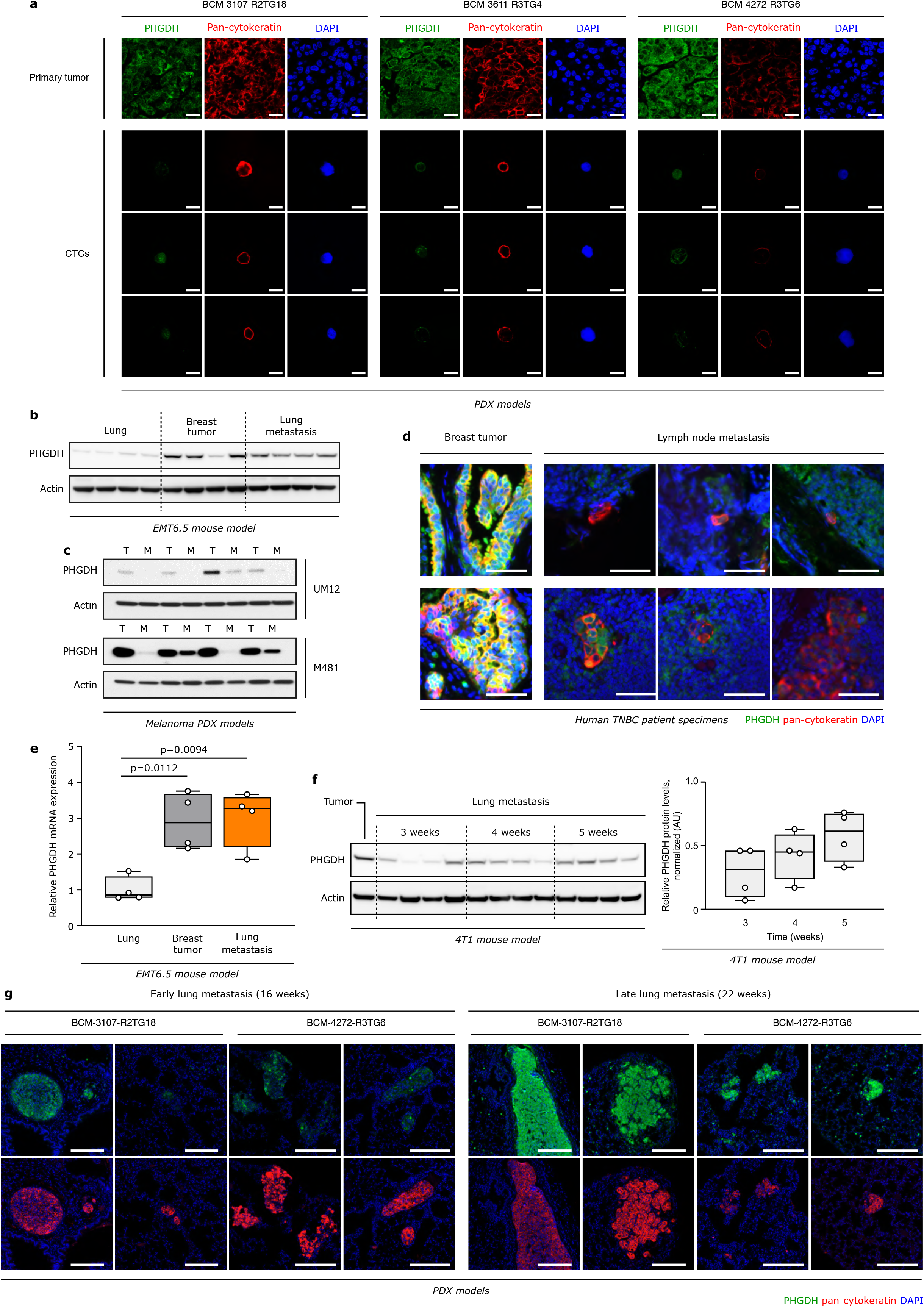
Circulating tumor cells and early metastatic lesions exhibit low PHGDH expression. **a.** Representative images of the expression levels of PHGDH protein in circulating tumor cells (CTCs) compared to the respective primary tumors from orthotopic (m.f.p.) TNBC PDX models, assessed by immunohistochemistry. Green, PHGDH; red, pan-cytokeratin tumor marker; blue, DAPI nuclear staining. Scale bar 25 μm. **b.** Western blot analysis of Phgdh in lungs, primary breast tumors and lung metastases from orthotopic (m.f.p.) EMT6.5 mouse model (n=4). **c.** Western blot analysis of Phgdh in primary tumors and lliver metastases from two different orthotopic (subcutaneous) melanoma PDX models (n=4). **d.** Representative images of the expression levels of PHGDH protein in lymph node metastases and matching primary breast tumors from TNBC patients, assessed by immunohistochemistry. Green, PHGDH; red, pan-cytokeratin tumor marker; blue, DAPI nuclear staining. Scale bar 50 μm. **e.** Relative change in *Phgdh* gene expression in lungs, primary breast tumors and lung metastases from orthotopic (m.f.p.) EMT6.5 mouse model (n=4). **f.** Western blot analysis of Phgdh in lung metastases from orthotopic (m.f.p.) 4T1 mouse model (n=4), at 3, 4 and 5 weeks after injection of the cancer cells. The box plot represents the relative quantification of the Phgdh band intensity over that of the housekeeper β-actin. **g.** Representative pictures of PHGDH protein expression in early (16 weeks) and late (22 weeks) lung metastases from orthotopic (m.f.p.) TNBC PDX models, assessed by immunohistochemistry. Green, PHGDH; red, pan-cytokeratin tumor marker; blue, DAPI nuclear staining. Scale bars 200 μm. The solid lines indicate the median, the boxes extend to the 25th and 75th percentiles, the whiskers span the smallest and the largest values. Unpaired t test with Welch’s correction, two-tailed.

**Extended Data Figure 2:**
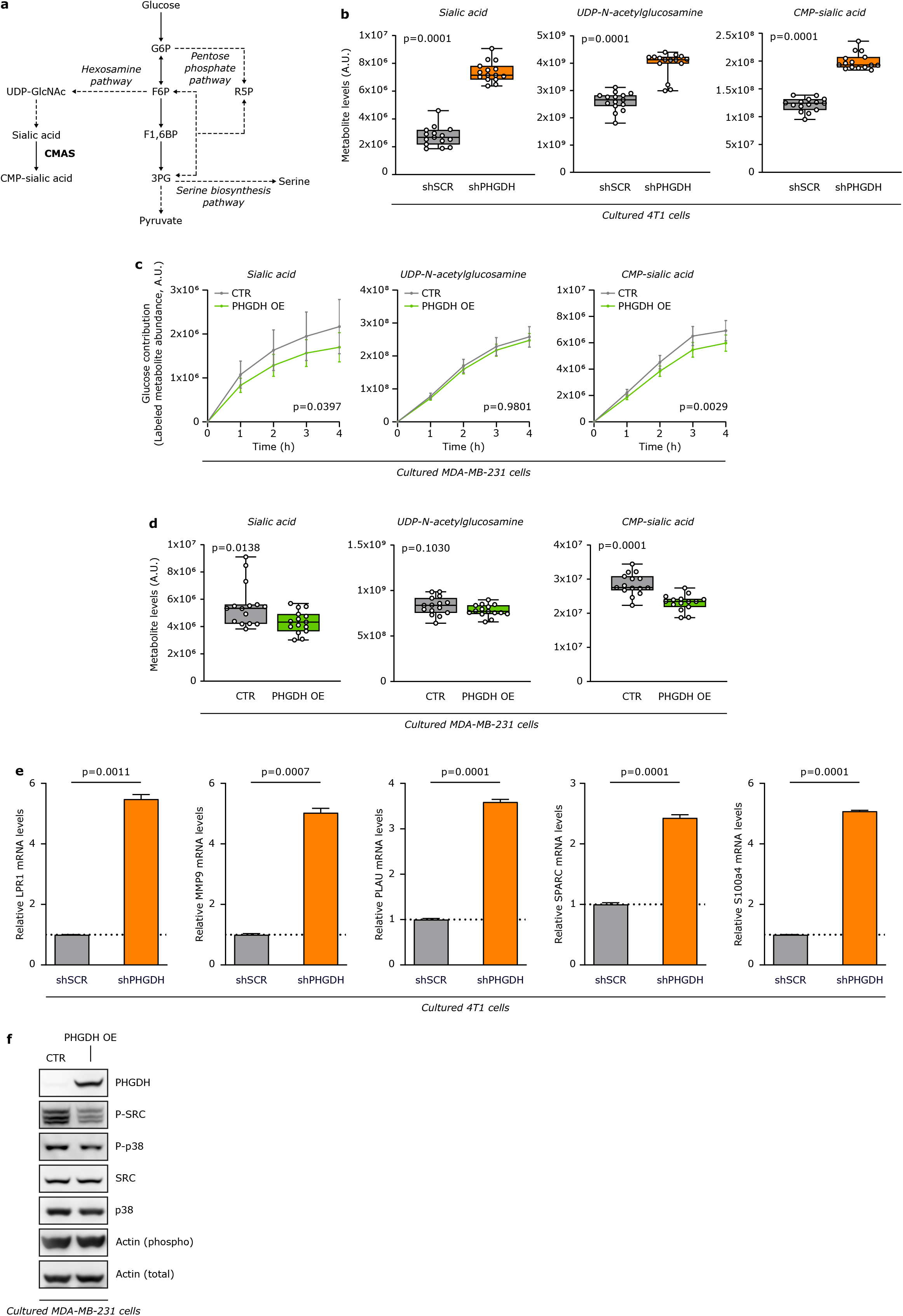
Low PHGDH expression promotes sialic acid synthesis and cellular programs downstream of sialic acid metabolism. **a.** Schematic representation of glycolysis and its branching metabolic pathways. Enzymes are depicted in bold, pathway names in italics. Solid lines represent single reactions, dashed lines recapitulate multiple reactions. **b.** Metabolite abundances of sialic acid, UDP-N-acetylglucosamine (UDP-GlcNAc) and CMP-sialic acid upon *Phgdh* knock-down in 4T1 cells. The solid lines indicate the median, the boxes extend to the 25th and 75th percentiles, the whiskers span the smallest and the largest values. N=15. Unpaired t test with Welch’s correction, two-tailed. **c.** Dynamic labeling of MDA-MB-231 cells, showing ^13^C glucose incorporation into sialic acid, UDP-N-acetylglucosamine and CMP-sialic acid upon PHGDH overexpression (PHGDH OE). Two-way ANOVA. Error bars represent s.d. from mean. **d.** Metabolite abundances of sialic acid, UDP-N-acetylglucosamine and CMP-sialic acid upon PHGDH overexpression (PHGDH OE) in MDA-MB-231 cells. The solid lines indicate the median, the boxes extend to the 25th and 75th percentiles, the whiskers span the smallest and the largest values. N=15. Unpaired t test with Welch’s correction, two-tailed. **e.** Relative change in *Lpr19*, *metalloproteinase 9* (*Mmp9*), *Plau*, *Sparc* and *S100a4* gene expression upon *Phgdh* knock-down in 4T1 cells. Error bars represent s.d. from mean. Unpaired t test with Welch’s correction, two-tailed. **f.** Phosphorylation of proto-oncogene tyrosine-protein kinase Src (c-SRC) and p38 mitogen-activated protein kinase upon PHGDH overexpression in MDA-MB-231 cells. N=3 unless otherwise stated. For western blot and expression data all experiments were performed in triplicate, one representative experiment is shown.

## RESOURCE AVAILABILITY

### Lead Contact

Further requests for resources should be directed to the lead contact, Sarah-Maria Fendt.

### Materials Availability

This study did not generate new unique reagents, except of genetically manipulated cell lines based on commercially available constructs. Reagents generated in this study will be made available on request through the lead author or the collaboration partner that generated the resource, but we may require a payment and/or a completed Materials Transfer Agreement if there is potential for commercial application.

### Data Availability

The datasets generated during and/or analyzed during the current study are available from the corresponding author on reasonable request.

### Code Availability

All the custom code used in the study is available from the corresponding author on reasonable request.

## METHODS

### Cell Culture

Human HEK293T epithelial cells, MDA-MB-231 breast adenocarcinoma cells and murine 4T1 mammary gland cancer cells were obtained from ATCC. Murine EMT6.5 mammary gland cancer cells were provided by R. Anderson (Peter MacCallum Cancer Center). 4T1 cells with CMAS knock-out (4T1 CMAS KO) and the respective control cells (4T1 CTR) were previously described and characterized^32^. HEK293T and MDA-MB-231 cells were cultured in high glucose (4.5 g/L) Dulbecco’s modified Eagle’s medium (DMEM) (Life Technologies) supplemented with 10% heat-inactivated fetal bovine serum (Life Technologies), 1% penicillin (final concentration of 50 U/mL) (Life Technologies) and 1% streptomycin (final concentration of 50μg/mL) (Life Technologies). 4T1 and EMT6.5 cells were cultured in Roswell Park Memorial Institute (RPMI) 1640 Medium (Life Technologies) supplemented with 10% heat-inactivated fetal bovine serum (Life Technologies), 1% penicillin (final concentration of 50 U/mL) (Life technologies) and 1% streptomycin (final concentration of 50μg/mL) (Life Technologies). The heat-inactivation of the fetal bovine serum was performed at 55°C for 45 minutes. Puromycin dihydrochloride (Life Technologies) and Hygromycin B (Life Technologies) were added to the growth medium for selection of knock-down and overexpression cell lines, respectively. All cell lines were confirmed to be mycoplasma free, based on routine testing with MycoAlert Mycoplasma Detection Kit (Lonza).

Cell-permeable α-ketoglutarate (dimethyl alpha-ketoglutarate, Sigma-Aldrich) was used at a concentration of 1 mM. The PHGDH inhibitor PH-755 was obtained from RAZE Therapeutics and used at a final concentration of 1 μM.

### Knock-down and overexpression strategies

PHGDH knock-down #1 in 4T1 cells was generated using the plasmid pLKO-shRNA2 vector expressing the shRNA against *Phgdh* obtained from the Peter Carmeliet’s lab (VIB/KU Leuven)^55^. The plasmids pLKO expressing the shRNA sequences for PHGDH knock-down #2 was obtained from BCCM (Belgian co-ordinated collections of micro-organisms). The same plasmid expressing a non-targeting shRNA sequence was used as control. Lentiviral particles were produced in HEK293T cells. Transduction of 4T1 cells was performed overnight and the medium was replaced the next day. Cells were selected with puromycin (2 μg/mL for 4T1 cells and 1 μg/mL for EMT6.5, 4T1 CMAS KO and 4T1 CTR cells). The overexpression of PHGDH in MDA-MB-231 cells was achieved using the plasmid pLHCX expressing the PHGDH cDNA obtained from the lab of Prof. Matthew Vander Heiden (MIT). The empty plasmid pLHCX was used as a control. Retroviral particles were produced in HEK293T cells. Transduction of MDA-MB-231 cells was performed overnight with freshly prepared virus and the medium was replaced the next day. Cells were selected with hygromycin (400 μg/mL). Polyclonal cells were selected for 1–2 weeks. Knock-down and overexpression of PHGDH were validated by western blot analysis (**Figure 4k, 4l, 4m, Extended Data Figure 3f**). 4T1 shSCR mTurquoise and 4T1 shPHGDH Dendra were obtained by using the plasmids pLenti-SFFV-H2B-Tq and pLKO-UBC-H2B-Dendra, respectively. Lentiviral particles were produced in HEK293T cells. Transduction of 4T1 cells was performed overnight and the medium was replaced the next day. mTurquoise- and Dendra-positive cells were FACS-sorted.

### Protein extraction and western blot analysis

Cells were collected in dPBS and lysed in RIPA lysis and extraction buffer (Thermo Scientific, 89901) supplemented with protease (Merck Sigma, 5892970001) and phosphatase (Merck Sigma, 4906845001) inhibitors. Frozen tissue samples were placed in 1.5 mL tubes with sufficient amount of RIPA buffer and grinded with a tissue lyser to help the lysis of the tissue.

Extracted proteins were quantified using a Pierce BCA Protein Assay Kit (Thermo Scientific, 23225). Subsequently, 10-40 μg of proteins were loaded on a precast gel NuPAGE Novex 4-12% Bis-Tris (Thermo Scientific, NP0336BOX). Proteins were then transferred onto a nitrocellulose membrane using an iBlot2 dry blotting system with iBlot2 transfer stacks (Thermo Scientific, IB301031). Membranes were incubated for 1h at room temperature in a blocking solution of 5% milk in TRIS Buffer Saline 0.05% Tween (TBS-T). Subsequently, membranes were incubated over night at 4 °C with primary antibodies against Phospho-SRC (Y416) (Cell Signaling Technology, 6943), SRC (Cell Signaling Technology, 2123), Phospho-p38 kinase (T180/Y182) (Cell Signaling Technology, 4511), p38 kinase (Cell Signaling Technology), β-actin (Merck Sigma, A5441), PHGDH (Merck Sigma, HPA021241), PSAT1 (Bio-techne, H00029968-A01), PFKL (Abcam, EPR11904), PFKM (Abcam, EPR10734(B)), PFKP (Abcam, EPR17314), or ALDOA (Novus Biologicals, NBP1-87488). All primary antibodies were used in a 1:1,000 dilution in 5% bovine serum albumin in TBS-T, except for PHGDH (1:3,000 dilution), β-Actin (1:10,000 dilution) and ALDOA (1:500 dilution). The following day, membranes were incubated with HRP-linked secondary antibodies anti rabbit (Cell Signaling Technology, 7074S) or mouse (Cell Signaling Technology, 7076S) used 1:4,000 in 5% milk in TBS-T. Bound antibodies were visualized using SuperSignal West Femto Maximum Sensitivity Substrate (Thermo Scientific, 34095) or SuperSignal West Pico PLUS Chemiluminescent Substrate (Thermo Scientific, 34580). Images were acquired using an ImageQuant LAS 4000 (GE Healthcare). Images were quantified using the software ImageQuant TL (GE Healthcare). The signals were normalized on the loading control β-Actin.

### Co-immunoprecipitation (IP)

Cells were collected in dPBS and lysed in CoIP lysis buffer (10 mM Tris-HCl pH 7.6, 140 mM NaCl, 5 mM EDTA, 0.5% Nonidet P-40) supplemented with protease (Merck Sigma, 5892970001) and phosphatase (Merck Sigma, 4906845001) inhibitors. Whole cell extracts (500 μg per IP) were immunoprecipitated for 1h using a PHGDH-specific antibody (Merck Sigma, HPA021241) complexed with Dynabeads Protein A (Life Technologies). After three washes with CoIP buffer, the isolated immunocomplexes were subjected to western blot analysis as described above.

### RNA isolation and RT-PCR

Total RNA was isolated with TRI reagent (Life Technologies). Quality and quantity of the isolated RNA was measured with a NanoDrop One Microvolume UV-Vis Spectrophotometer (Thermo Scientific). RNA was reverse transcribed into cDNA using a qScript cDNA Synthesis Kit (Quantabio). The relative levels of transcripts compared to the control RPL19 were determined by qPCR using PerfeCTa SYBR Green SuperMix, Low ROX (Quantabio) and specific primers on a 7500 Fast Real Time PCR System (Applied Biosystems, Life Technologies). Amplification was performed at 95 °C for 10 min, followed by 40 cycles of 15 s at 95 °C and 1 min at 60 °C.

The primer list is provided in **Extended Data Table 5**.

### Bulk RNAseq analysis

RNA from freshly collected cells was extracted with TRI reagent (Life Technologies). Quality and quantity of the isolated RNA was measured with a NanoDrop One Microvolume UV-Vis Spectrophotometer (Thermo Scientific). Total RNA of 4T1 cells with either knockdown of PHGDH or control (3 replicates each) was collected as described above. RNAseq libraries were prepared from 1 μg of total RNA per sample using the KAPA Stranded mRNA Sequencing Kit (Roche). In short, poly-A containing mRNA was purified from total RNA using oligo(dT) magnetic beads and fragmented into 200–500 bp pieces using divalent cations at 94°C for 8 min. The cleaved RNA fragments were copied into first strand cDNA. After second strand cDNA synthesis, fragments were A-tailed and indexed adapters were ligated. The products were purified and enriched by PCR to create the final cDNA library. After quantification with qPCR, the resulting libraries were sequenced on a HiSeq4000 (Illumina) using a flow cell generating 1×50bp single-end reads.

Resulting reads were trimmed for adaptors and low quality basecalls using Trim Galore! (v0.5.0), mapped to the mm10 genome assembly using HISAT2 (v2.1.0) (PMID: 25751142) and quantified using default settings with featureCounts (v1.6.4) (PMID: 14668230) as implemented in the nf-core rnaseq pipeline (v1.3). Differential expression of knockdown vs control was performed using the DESeq2 R package (v1.26.0). Resulting log2(fold change) were ranked and used as input for a gene set enrichment analysis using a GSEA graphical tool (https://software.broadinstitute.org/gsea/index.jsp) using all mouse biological product GO terms, except the electronically inferred terms (downloaded from http://download.baderlab.org/EM_Genesets/current_release/Mouse/symbol/) or a mouse ortholog version of the human Molecular Signature database (downloaded from http://download.baderlab.org/EM_Genesets/July_05_2019/Mouse/symbol/Pathways/Mouse_Human_MSigdb_July_05_2019_symbol.gmt) as gene sets in a preranked analysis. Only gene sets containing between 15 and 500 genes were retained and the number of permutations was set at 1,000.

### Orthotopic mouse models

All animal experiments were approved by the local authorities in compliance with all relevant ethical regulations. For injection models, mice were randomized before injection of cancer cells. All samples were analyzed blinded. Sample size was determined using power calculations with B = 0.8 and P < 0.05 based on preliminary data and in compliance with the 3R system: Replacement, Reduction, Refinement.

Mice were housed in filter top cages and IVC cages. Housing and experimental animal procedures were approved by the Institutional Animal Care and Research Advisory Committee of KU Leuven, Belgium. The animal study complies with ethical regulations and was approved by the KU Leuven ethics committee.

Six-week-old female BALB/c (Envigo) mice were inoculated in the mammary fat pad with 1×10^6^ cancer cells, in 50 μL PBS using a 29G syringe. Mice were euthanized 18-21 days after cell injection, except for experiments in **Extended Data Fig. 1e**, in which tumors were resected one week post-injection and mice were euthanized 3, 4 or 5 weeks post-injection. At the end of the experiment, primary tumors were dissected and weighted. Humane end points were determined as follows: tumor size of 1.8 cm^3^, loss of ability to ambulate, labored respiration, surgical infection or weight loss over 10% of initial body weight. Mice were monitored and upon detection of one of the previous mentioned symptoms, the animal was euthanized.

At the end of every *in vivo* experiment, mice were sacrificed by injecting approximately 50 μL of a 60 mg/mL Dolethal (pentobarbital sodium) solution (Vetoquinol). Primary tumors, lung metastases and lung healthy tissue were dissected and washed in ice-cold saline, placed into pre-labelled bags and frozen using a liquid nitrogen-cooled Biosqueezer (Biospec Products). The bags were then placed in liquid nitrogen until all collections were finished and finally stored at −80 °C until further processing.

### Patient-derived xenografts (PDX) tumor models

All animal experiments were performed under IACUC approved protocols at Baylor College of Medicine (Houston, TX). Triple negative breast cancer PDX models^56^ were screened via IHC for expression of PHGDH in the tumor, lung, liver, and brain. Three models (BCM-3107, BCM-3611 and BCM-4272) were chosen based on the positivity for PHGDH in the primary tumor. Fresh PDX tumor tissue fragments were transplanted into the cleared fourth fat-pad (right abdominal) of four-week-old SCID/Beige mice^57^.

In a first cohort of 6 mice, when tumors reached a size of ~500mm^3^ (length × width^2^ × 0.5), the animals were sacrificed and blood was collected and processed for circulating tumor cell (CTC) isolation. A portion of the tumor was fixed overnight in 10% neutral buffered formalin and placed in 70% ethanol until paraffin embedding (FFPE) and the remainder was snap frozen in liquid nitrogen. In a second cohort of 4 mice, when tumors reached a size of ~500mm^3^ (length × width^2^ × 0.5), tumor resection surgeries were performed to remove the primary tumor. Two months after tumor resection, animals were sacrificed.

At the terminal endpoint for each cohort, lungs were collected fixed overnight in 10% neutral buffered formalin and placed in 70% ethanol until paraffin embedding (FFPE).

Melanoma specimens were obtained with informed consent from all patients according to protocols approved by the Institutional Review Board (IRB) of the University of Michigan Medical School (IRBMED approvals HUM00050754 and HUM00050085^58^) and the University of Texas Southwestern Medical Center (IRB approval 102010-051). Single-cell suspensions were obtained by dissociating tumors mechanically with a scalpel on ice. Cells were filtered through a 40-μm cell strainer to remove clumps.

All mouse experiments complied with all relevant ethical regulations and were performed according to protocols approved by the Institutional Animal Care and Use Committee at the University of Texas Southwestern Medical Center (protocol 2016-101360). For all experiments, the maximum permitted tumor diameter was 2.5 cm and this limit was not exceeded in any experiment. For all experiments, mice were kept on normal chow and fed ad-libitum. Melanoma cell suspensions were prepared for injection in staining medium (L15 medium containing bovine serum albumin (1 mg ml−1), 1% penicillin–streptomycin and 10 mM HEPES (pH 7.4) with 25% high-protein Matrigel (354248; BD Biosciences)).

Patient-derived melanomas were transplanted into 4-to-8-week-old male and female NOD.CB17-Prkdc^scid^ Il2rg^tm1Wjl^/SzJ (NSG) mice. Subcutaneous injections were performed in the right flank of mice in a final volume of 50 μl using 100 cells per injection for human melanoma cells. Subcutaneous tumor diameters were measured weekly with calipers until any tumor in the mouse cohort reached 2.5 cm in its largest diameter. At that point, all mice in the cohort were euthanized, per approved protocol, for collection of subcutaneous tumors and macrometastatic nodules.

### Collection of Circulating Tumor Cells

Whole blood collected from the PDX-bearing mice was diluted 1:40 in PBS. Diluted blood was processed using VTX-1 microscale vortices technology (Vortex BioSciences) to collect live circulating tumor cells directly onto microscope slides containing a FlexiPERM ring. Slides were placed in a 4-well dish, spun, and placed in a 37°C incubator for one hour to allow cells to attach. Cells were then fixed with 4% PFA. Two washes with PBS were used to remove the PFA and the slides were allowed to air dry prior to staining.

### Histopathology and immunohistochemistry (IHC)

Immunohistochemical analyses were performed on human breast cancer tissues from the tissue archives of the Institute of Pathology of the LMU Munich. Grade 2/3 invasive ductal carcinomas of the breast, not otherwise specified (NOS), and treated by primary surgical resection between 1988 and 2006 were investigated. Samples were selected for triple-negative breast cancers based on routine immunophenotypical profiling for hormone receptors (ER/PR) and Her2-neu expression. Tumor samples had been anonymized and analyzed according to the local ethics committee regulations (19-914 KB). Tissue samples from representative lesions were collected and fixed in 4% paraformaldehyde for 24 hours and then processed for paraffin embedding (HistoStar™ Embedding Workstation). Sections of 4 μm of thickness obtained from the paraffin-embedded tissues (Thermo Scientific Microm HM355S microtome) were mounted on Superfrost™ Plus Adhesion slides (Thermo Scientific) and routinely stained with hematoxylin and eosin (H&E, Diapath #C0302 and #C0362) for histopathological examination.

#### Hematoxylin and eosin staining

Hematoxylin and eosin staining was used to identify the cancerous lesions in the lung of the different mouse models. During dissection, the lungs were gently infused via the trachea with 10% neutral buffered formalin. Next, samples were embedded in paraffin and sliced in 7 μm thick sections that were stained with hematoxylin and eosin. Images were acquired on a Zeiss Axio Scan.Z1 using a x20 objective and ZEN 2 software and analyzed using the ZEN Blue software (Zeiss). In order to focus on the early stages of the metastatic process, only mice bearing lung metastases ≤0.1 mm^2^ (4T1) or ≤0.4 mm^2^ (EMT6.5) were analyzed. The rate of metastatic progression was calculated over a 3-day period, comparing the average metastatic burden (MB) in the lung (measured as total metastasis area) of 10 mice per group at 18 and 21 days post orthotopic injection with either 4T1 shSCR or 4T1 shPHGDH cells. The following formula was used: rate of metastatic progression = (ln(MB_21days_) - ln(MB_18days_) / ln(2) × time.

#### Immunofluorescence with and without tyramide signal amplification

Tissue sections of 4 μm were deparaffinized and hydrated in distilled water, followed by 23 min of heat-induced epitope retrieval (HIER) in AR6 buffer (Perkin Elmer, AR6001KT) using the 2100 Antigen Retriever (Aptum Biologics Ltd). After a cooldown of 15 min in milliQ water the endogenous peroxidase activity of the samples was blocked by a 20 min incubation in 0.3 % hydrogen peroxide in methanol. The tissue was then blocked for 30 min using TNB blocking buffer (0.1 M TRIS-HCl, pH 7.5; 0.15 M NaCl; 0.5% TSA Blocking Reagent (PerkinElmer, FP1012 or FP1020)). Pan-cytokeratin (panCK) was detected by incubating the tissue with anti-panCK antibody (mouse, Agilent, M351501-2, 1:100 in TNB blocking buffer) for 30 min. Antibody detection was followed by 45 min of incubation with goat anti-mouse-biotin (DAKO, E0433, 1:200 in TNB blocking buffer) and subsequent incubation with Streptavidin-HRP Conjugate (PerkinElmer, NEL750001EA, 1:100 in TNB blocking buffer). Signal detection was performed by 8 min incubation with the PerkinElmer TSA Plus Cyanine 3 kit (PerkinElmer, NEL744E001KT, 1:50 in amplification buffer from the Cyanine 3 kit). For the detection of PHGDH, the tissue was first blocked for 30 min using a blocking buffer (TBS with 1% BSA (VWR, 22013, Bovine Serum Albumin (BSA), fraction V, Biotium (50 g))) with 10% normal goat serum (Invitrogen, 10000C) and then incubated with anti-PHGDH antibody (rabbit, Merck Sigma, HPA021241, 1:3,000 in TNB blocking buffer) for 30 min. The PHGDH antibody was visualized by incubating the sections with goat anti-rabbit Alexa Fluor 647 antibody (Goat anti-Rabbit IgG (H+L) Highly Cross-Adsorbed Secondary Antibody Alexa Fluor 647, Lifetech, A21245, 1:200 in TNB blocking buffer) for 45 min followed by DAPI staining for 5 min (Spectral DAPI, Akoya Biosciences, FP1490, 2 drops per 1 ml TBST). The slides were then mounted using ProLong Diamond Antifade Mountant (Thermo Fisher, P36961).

For triple stainings for PHGDH, panCK and PHH3, the following antibodies were used: anti-panCK (mouse, 1:1,000; DAKO, M3515, Clone AE1/AE3), anti-PHH3 (rabbit, Cell Signaling Technology, #9701S), and anti-PHGDH antibody (rabbit, 1:10,000 in TNB blocking buffer, Merck Sigma, HPA021241). The PerkinElmer Opal 4-Color Manual IHC Kit (PerkinElmer/Akoya, NEL810001KT) was used for the tyramide signal amplification according to the manufacturer’s protocol. For introduction of the secondary-HRP the Envision+/HRP goat anti-Rabbit (Dako Envision+ Single Reagents, HRP, Rabbit, Code K4003) was used for antibody raised in rabbit (PH3 and PHGDH) and the OPAL Polymer HRP Ms+Rb (Akoya/Perkin Elmer, ARH1001EA) was used for the antibody raised in mouse (panCK). The slides were then mounted using ProLong Diamond Antifade Mountant (Thermo Fisher, P36961). The various proteins were detected by first using the OPAL 570 (panCK), then OPAL 690 (PHH3), and at last OPAL 520 (PHGDH) reagents according to the manufacturer’s protocol.

#### Microscope image acquisition and image processing

Images were acquired on a Zeiss Axio Scan.Z1 using a x20 objective and ZEN 2 software. For exporting images the ZEN 2 software (Zeiss) and the software package QuPath (Version: 0.1.2,^59^) were used. QuPath was also used for automatic cell detection using the DAPI channel and for subsequent creation of a detection classifier using all 55 given parameters resulting in the classification of panCK-positive cells within the whole slide.

### Imaging Mass Cytometry (IMC)

#### Tissue preparation and staining

Carrier-free antibodies were labelled with stable metal isotopes (**Extended Data Table 6**) using the MaxPar labeling kit (Fluidigm) following the manufacturer’s instructions. Antibodies were quantified photometrically and stored at 4 °C.

Formalin-Fixed Paraffin-Embedded (FFPE) tumors were fixed in 10% NBF overnight and embedded in paraffin. 5um sections were baked at 60 °C for 2 hours before dewaxing in xylene for 20 min. Samples were then rehydrated in descending grades of ethanol (100%, 95%, 89%, 70%) 5 min each. Antigen retrieval was performed in Tris-EDTA HIER buffer pH 9.2 at 96 °C for 30 min. Samples were cooled at room temperature before blocking. FFPE sections were blocked in TBS/0.3% Triton X-100/3% BSA for 1 hour. Samples were incubated overnight at 4 °C in primary antibody panel at 5 ug/mL each diluted in TBS/0.1% Triton X-100/1% BSA in a humid chamber. The antibody panel used is described in **Extended Data Table 6.** Samples were washed three times in TBS and once in milliQ water before nuclei staining with MaxPar Intercalator-Ir (Fluidigm) diluted 1:400 in TBS for 15 min. Samples were washed once in milliQ water for 5 min and air dried before imaging.

#### Data acquisition

IMC acquisitions were performed on a Fluidigm Hyperion instrument connected to a Fluidigm Helios mass cytometer. Laser frequency was 200 Hz with a raster scanning resolution of 1 μm (XY). The abundances of each isotope per pixel were measured and converted in image by the Fluidigm Cytof software.

#### Data analysis – Single cell segmentation and quantification

IMC images were segmented into single cells following using the packages imctools, Ilastik 1.2.2^60^ and CellProfiler 3.1.5^61^ as described in the IMCSegmentationPipeline^62^ available at https://github.com/BodenmillerGroup/ImcSegmentationPipeline. In brief, imctools was used to convert MCD and txt files obtained from imaging into TIFF format. Supervised pixel classification was done with Ilastik using a combination of nuclear, cytoplasm and cell membrane markers to generate probability maps of single cells (nuclei, cytoplasm/membrane and background). Probability maps were then used to generate a segmentation mask using CellProfiler. Finally, segmentation masks and TIFF images for all channels of interest were overlaid to filter out outlier pixels and extract single-cell measurements for each channel.

#### Data analysis – Clustering and correlation analysis

Single-cell mean intensities for each channel were analyzed using the R package Seurat 3.2 (http://satijalab.org/seurat/). Cell outliers corresponding to hot pixels were filtered out. The remaining cells were scaled and clustered using a graph-based approach. For high-dimensional clustering all measured channels were used (**Extended Data Table 6**). For visualization, high-dimensional single-cell data were reduced to two dimensions using UMAP. Tumor cells were selected by a negative selection of clusters corresponding to immune cells (CD45+), stromal cells (aSMA+) and endothelial cells (CD31+ and/or CD34+). Data distribution was tested for normality. Because all variables were not normally distributed, correlation of expression between PHGDH and Ki67 in tumor cells was investigated with the nonparametric Spearman rank correlation. Differences were considered significant at p<0.05.

### Time-lapse intravital microscopy

NOD-scid Il2ry^null^B2m^null^ (NSG) mice were obtained from Jackson Laboratories. All animal experiments were approved by the Animal Welfare Committee of the Netherland Cancer Institute (NKI), in accordance with national guidelines. All animals were maintained in the animal department of the NKI, housed in individually ventilated cage (IVC) systems under specific pathogen-free conditions and received food and water *ad libitum*. Sample size was determined using power calculations with B = 0.8 and P < 0.05 based on preliminary data and in compliance with the 3R system: Replacement, Reduction, Refinement.

#### Time-lapse intravital imaging of primary tumor

4T1 shCRTL-mTq 4T1 shPDGDH-Dendra cells (50,000 cells per injection) were mixed in 100 μl Matrigel (Corning, cat. no. 356231) and injected into the 4th mammary fat pad of recipient NSG mice. Experiments were performed on 8 NSG mice and time-laps intravital imaging was performed on tumors of about 200 mm^3^. During the entire procedure mice were sedated using isoflurane inhalation anesthesia (~1.0% isoflurane/ compressed air mixture) and received 200 μl sterile PBS by subcutaneous injection before the surgery. The tumors were surgically exposed and mice were then placed in a custom designed imaging box on the microscope while kept under constant anesthesia, with the imaging box and the microscope adjusted to 34.5 °C using a climate chamber. Intravital images were acquired using an inverted Leica SP8 Dive system equipped with a MaiTai eHP DeepSee laser (Spectra-Physics) and Insight X3 (Spectra-Physics) and 4 HyD-RLD detectors. mTurquosie and Dendra were simultaneously excited at 820 nm (Mai Tai) and 960 nm (Insight X3), respectively, and singles including second harmonic generation (Collagen I, stroma) were detected using the hybrid detector. Different positions within the tumor were acquired as three-dimensional tile scans with a 5 μm Z-steps. Images were collected every 30 minutes for a period of 7 to 9 h during which the mouse was kept sedated and alive, constantly hydrated with subcutaneous infusion of glucose and electrolytes (NutriFlex special 70/240, Braun, 100 μl/h). All images were collected at 12 bit and acquired with a 25x water immersion objective with a free working distance of 2.40 mm (HC FLUOTAR L 25x/0.95 W VISIR 0.17).

#### Post-processing and analysis of time-lapse intravital microscopy

The time-lapse three-dimensional positions were corrected for XYZ-drift using the Huygens Object Stabilizer module (Scientific Volume Imaging). The correctness of the XY correction was visually inspected and manually adjusted using ImageJ if required. Migration of randomly picked individual cells was analyzed using the MTrack2 plugin in ImageJ (http://imagej.net/MTrack2). Only cells that could be followed over a minimal period of 4 h were included in the analysis. Cells were defined as being migratory when the mean displacement per hour was greater than 4 μm. For track length analysis, only cells which could be followed over the whole duration of time-lapse imaging were included in the analysis.

### Transwell migration assay

4T1 mTq and EMT6.5 mTq cells, both shSCR and shPHGDH, were seeded in ClearView 96-well Cell Migration Plates (Essen BioScience) at 4×10^3^ cells/well. Where indicated in the figure legends, transwells were pre-coated with a confluent monolayer of either primary human umbilical vein endothelial cells (HUVECs) or primary lymphatic endothelial cells (LECs). Time-lapse monitoring of cancer cell migration was performed over a period of 48 h using the IncuCyte ZOOM Live-Cell Imaging Instrument equipped with the IncuCyte Chemotaxis Cell Migration Software Module (Essen BioScience). The number of migrated cancer cells, defined as mTq-positive cells, was determined in each well by analyzing the time-lapse data with the Incucyte Zoom software (Essen BioScience).

### *In vitro* invasion assay

Cancer cells were tested for *in vitro* invasion either as loose cells (4T1, EMT6.5 and MDA-MB-231) or spheroids (4T1). For the invasion assay with loose cells, 50,000 cells were embedded in a 50:50 mix of growth factor-reduced Matrigel (BD Biosciences) and collagen I (Life Technologies) and seeded onto 35 mm glass bottom culture dishes (MatTek). Cells were allowed to invade for 48 h (EMT6.5) or 72 h (4T1 and MDA-MB-231), then they were stained with calcein green (Life Technologies) for 1h, washed with PBS and immediately imaged. For the invasion assay with spheroids, 6,000 cancer cells were grown in hanging drops in full medium for 72 h, then the spheroids were collected, embedded in a 50:50 mix of growth factor-reduced Matrigel (BD Biosciences) and collagen I (Life Technologies) and seeded onto 35 mm glass bottom culture dishes (MatTek). Cells were allowed to invade for 72h, then they were stained with calcein green (Life Technologies) for 1h, washed with PBS and immediately imaged. Imaging was performed on a Leica TCS SP8 X confocal microscope equipped with a White Light Laser and a HCX PL APO CS 10x/0.40 DRY objective. Images were acquired as three-dimensional scans with 10 μm Z-steps and processed with LAS X software (Leica) to obtain maximum projection images. Quantification of invasive area and invasive distance was performed on the maximum projection images using the FIJI^63^ distribution of ImageJ2^64^.

### Metabolite measurements

All labeling experiments were performed in media with 10% dialyzed serum for the indicated time points. Metabolites for the subsequent mass spectrometry analysis were prepared by quenching the cells in liquid nitrogen followed by a cold two-phase methanol-water-chloroform extraction^65^. Phase separation was achieved by centrifugation at 4 °C. The methanol-water phase containing polar metabolites was separated and dried using a vacuum concentrator. Dried metabolite samples were stored at −80 °C. The protein interphase was also dried down and dissolved in 200 μl 0.2 M potassium hydroxide, and the protein concentration was then quantified using the Pierce BCA Protein Assay Kit (Thermo Fisher).

#### Gas chromatography-mass spectrometry (GC-MS)

Serine and glycine were measured with gas chromatography-mass spectrometry. Polar metabolites were derivatized for 90 min at 37 °C with 20 μl of 20 mg per ml methoxyamine (Merck Sigma, 226904) in pyridine (Merck Sigma, 270970). Subsequently, 15 μL of N-(tert-butyldimethylsilyl)-N-methyl-trifluoroacetamide, with 1 % tert-butyldimethylchlorosilane were added to 7.5 μL of each derivative and incubated for 60 min at 60 °C (Merck Sigma, 375934)^65^. Isotopologue distributions and metabolite concentrations were measured with a 7890 A GC system (Agilent Technologies) combined with a 5975C Inert MS system (Agilent Technologies). 1 μl of sample was injected into a DB35MS column in split mode (ratio 1 to 3) using an inlet temperature of 270 °C. The carrier gas was helium with a flow rate of 1 ml/ min. Upon injection, the GC oven was set at 100 °C for 1 min and then increased to 105 °C at 2.5 °C/min and with a gradient of 2.5 °C/ min finally to 320 °C at 22 °C/min. The measurement of metabolites was performed under electron impact ionization at 70 eV using a selected-ion monitoring (SIM) mode. Isotopologue distributions were extracted from the raw ion chromatograms using a custom Matlab M-file, which applies consistent integration bounds and baseline correction to each ion^66^. In addition, we corrected for naturally occurring isotopes^67^. All labeling fractions were transformed into natural abundance-corrected mass distribution vectors (MDVs)^33^.

#### Liquid chromatography-mass spectrometry (LC-MS)

For the detection of metabolites by LC-MS, a Dionex UltiMate 3000 LC System (Thermo Scientific) with a thermal autosampler set at 4 °C, coupled to a Q Exactive Orbitrap mass spectrometer (Thermo Scientific) was used. Samples were resuspended in 50 μL of water and a volume of 10 μl of sample was injected on a C18 column (Acquity UPLC HSS T3 1.8 μm 2.1×100 mm). The separation of metabolites was achieved at 40 °C with a flow rate of 0.25 ml/min. A gradient was applied for 40 min (solvent A: 10mM Tributyl-Amine, 15 mM acetic acid – solvent B: Methanol) to separate the targeted metabolites (0 min: 0% B, 2 min: 0% B, 7 min: 37% B, 14 min: 41% B, 26 min: 100% B, 30 min: 100% B, 31 min: 0% B; 40 min: 0% B. The MS operated in negative full scan mode (m/z range: 70-1050 and 300-700 from 5 to 25 min) using a spray voltage of 4.9 kV, capillary temperature of 320 °C, sheath gas at 50.0, auxiliary gas at 10.0. Data was collected using the Xcalibur software (Thermo Scientific) and analyzed with Matlab for the correction of protein content and natural abundance, but also to determine the isotopomer distribution using the method developed by Fernandez et al, 1996^67^

#### Glycolytic flux

4T1 cells were cultured for 6 h in RPMI medium containing 0.4 μCi ml^−1^ [5-^3^H]D-glucose (Perkin Elmer) after which the supernatant was transferred into glass vials sealed with rubber stoppers. ^3^H2O was captured in hanging wells containing a Whatman paper soaked with H_2_O over a period of 48 h at 37 °C to reach saturation. The paper was then used for liquid scintillation counting (QuantaSmart V4 Perkin Elmer).

#### Glucose uptake

Glucose uptake was measured in 4T1 cells using the fluorescent glucose analog 2-deoxy-2-[(7-nitro-2,1,3-benzoxadiazol-4-yl)amino]-D-glucose (2-NBDG; Cayman Chemicals). Briefly, the cells were seeded in 96-well plates, cultured for 1 h in glucose- and FBS-free RPMI medium containing 200 μg/ml 2-NBDG and washed three times in PBS. Glucose uptake was assessed by spectrophotometric measurement of the fluorescence emitted by the cells at 550 nm. Fluorescence readings were normalized over cell count.

### Statistical analysis

Statistical data analysis was performed using GraphPad Prism version 8.0 (GraphPad Software) on n ≥ 3 biological replicates. Details on statistical tests and post-tests are presented in the figure legends. Sample size for all experiments was chosen empirically. Independent experiments were pooled and analyzed together whenever possible as detailed in figure legends. Data are presented as mean ± s.d., mean ± s.e.m. or median ± 95% Confidence Interval, as indicated in the figure legends.

